# Multi-Approach and Multi-Scale Model of CD4+ T Cells Predicts Switch-Like and Oscillatory Emergent Behaviors in Inflammatory Response to Infection

**DOI:** 10.1101/2020.02.26.964502

**Authors:** Kenneth Y. Wertheim, Bhanwar Lal Puniya, Alyssa La Fleur, Ab Rauf Shah, Matteo Barberis, Tomáš Helikar

## Abstract

Immune responses rely on a complex adaptive system in which the body and infections interact at multiple scales and in different compartments. We developed a modular model of CD4+ T cells which uses four modeling approaches to integrate processes taking place at three spatial scales in different tissues. In each cell, signal transduction and gene regulation are described by a logical model, metabolism by constraint-based models. Cell population dynamics are described by an agent-based model and systemic cytokine concentrations by ordinary differential equations. A Monte Carlo simulation algorithm allows information to flow efficiently between the four modules by separating the time scales. Such modularity improves computational performance and versatility, and facilitates data integration. Our technology helps capture emergent behaviors that arise from nonlinear dynamics interwoven across three scales. Multi-scale insights added to single-scale studies allowed us to identify switch-like and oscillatory behaviors of CD4+ T cells at the population level, which are both novel and immunologically important. We envision our model and the generic framework encompassing it to become the foundation of a more comprehensive model of the human immune system.

---

Immune responses mediated by CD4+ T cells involve complex interactions among immune cells and molecules. Resting CD4+ T cells are activated by antigen-presenting cells and cytokines, further differentiate, and secrete cytokines to act against pathogens and abnormal cells. They also recruit other immune cells to the sites of infection. Depending on the cytokine milieu, activated CD4+ T cells may differentiate into various phenotypes including T helper type 1 (Th1), T helper type 2 (Th2), T helper type 17 (Th17), and induced T regulatory cells (Tregs) [1]. To produce the energy and molecular precursors required to achieve a specific mixture of phenotypes, activated CD4+ T cells utilize certain signaling and metabolic pathways such as aerobic glycolysis [1,2]. To fully understand these complex interactions underlying the dynamics of CD4+ T cell immune response, we must integrate events taking place at various spatial, temporal, and organizational scales, such as immune cell proliferation, development, and migration; cell-cell and cell-molecule interactions; intracellular signaling; and intracellular metabolism.

Multi-scale modeling aims to integrate spatial, temporal, and organizational scales of biological systems. This strategy has been used extensively in immunology. For instance, multi-scale models have been developed to study infections and inflammatory processes [3], and the immune response to the *Helicobacter pylori* infection [4]. Such integration could be achieved by combining different modeling approaches, such as ordinary differential equations (ODEs) and partial differential equations (PDEs) for the chemical kinetics and transport of molecular species (in terms of concentrations) in and across different cells, organs or tissues; agent-based modeling (ABM) for cells and molecules interacting in heterogeneous spatial environments; and ODE/PDE-based, rule-based, logic-based, or constraint-based models for intracellular dynamics. Genome-Scale Metabolic Models (GSMMs) have been developed for human tissues and cells [5,6] and can be used to model whole-cell metabolism in a context-specific manner. Logic-based models can be used to model cell signaling and gene regulation where kinetic parameters are unavailable [7]. However, previous modeling studies did not consider all these scales, consider multiple compartments/tissues, or employ appropriate modeling approaches for their selected scales. Moreover, the resulting modeling frameworks are difficult to extend and re-use in different contexts.

Herein, we present a multi-approach and multi-scale modeling framework that can be used to model diverse immune responses at molecular, cellular, and systemic scales. We demonstrated its capabilities by modeling the dynamics of CD4+ T cells in response to influenza infections (real and hypothetical) in different cytokine milieus, considering heterogeneous populations of Th0, Th1, Th2, Th17, and Treg subtypes and 11 cytokines in three spatial compartments (an infection site or target organ, lymphoid tissues, and a circulatory system). For each subtype, we modeled the intracellular signaling and gene regulation network underpinning the cell’s proliferation, differentiation into that subtype, as well as the whole-cell metabolism of that subtype. Based on our simulations of CD4+ T cell responses to the hypothetical infections, we predicted novel emergent behaviors.

## Results

### Four different modeling approaches represent processes spanning three spatial scales in three compartments

The multi-scale model is divided into three compartments (Fig. 1). The first target organ compartment represents the site of initial insult (*e*.*g*., infection). Within the context of the validation and robustness studies presented below, pertaining to influenza infections, this compartment represents the lungs. The second compartment is an abstract representation of the lymphoid tissues connected to the target organ: a micro-environment where CD4+ T cells can be stimulated by antigens. In the case of influenza, this compartment represents the draining lymph nodes nearest the lungs. The final compartment is the circulatory system, which is an abstraction of the rest of the body. Each compartment is well-stirred with respect to both cells and cytokines. The CD4+ T cells in each compartment can sense and/or secrete 11 cytokines, and they are stimulated by a study-dependent and time-varying signal which represents the antigen load and the activity of the rest of the immune system. A CD4+ T cell is created as a naive cell with the Th0 phenotype and typically goes through three major stages: activation, expansion, and contraction (including memory formation and cell death). The behaviors and attributes of each cell are dynamic and depend on the outputs of the intracellular models of signal transduction, gene regulation, and metabolism. The processes related to signal transduction and gene regulation include the canonical pathways regulating the transdifferentiation of effector CD4+ T cells (interconversion among mixtures of Th0, Th1, Th2, Th17, and Treg phenotypes) [8], and events such as aerobic glycolysis, memory formation, and cell death. These pathways are detailed in the Supplementary Text. Although the metabolic models consider whole-cell metabolism at the genome-scale, different subsets of metabolic fluxes are used to model different parts of the cell cycle in the multi-scale model. For instance, the fluxes related to glutaminolysis are active during the S phase when DNA is synthesized.

**Figure 1:**
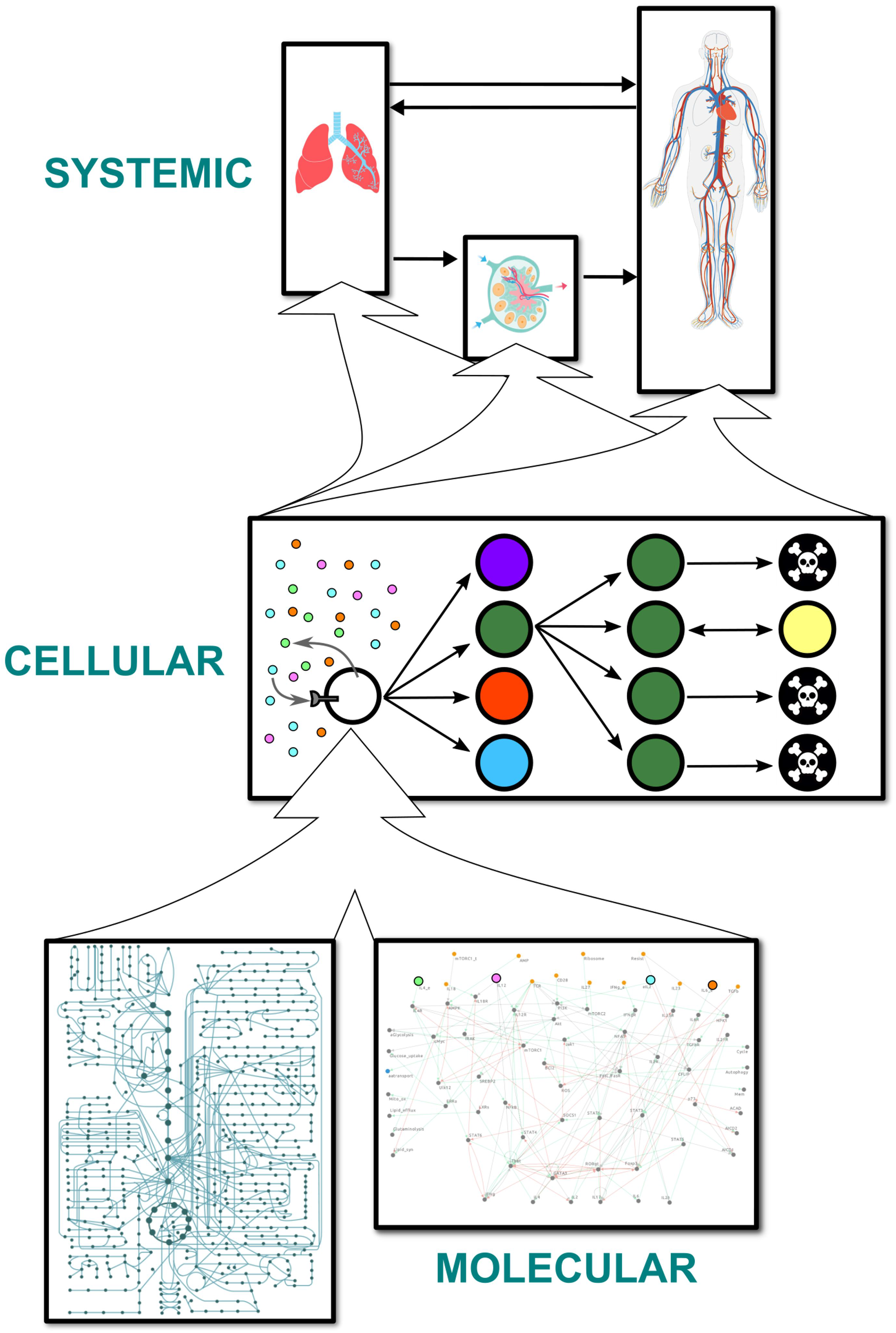
overall structure of the model. Schema representing the arrangement of models at three spatial scales. Cells are represented by discrete, autonomous agents that can differentiate into different subtypes, divide, and die. Each cell contains a constraint-based metabolic model for each phenotype and a logical model of signaling and gene regulatory pathways. The agents themselves reside in and travel between three isotropic compartments. Each compartment is associated with a set of cytokine concentrations, modeled by ordinary differential equations, that affect and are affected by the agents.

The aforementioned processes are represented using four different modeling frameworks. **At the systemic scale**, 11 sets of three coupled compartmental ordinary differential equations model the temporal dynamics of the 11 cytokine concentrations in the three compartments. Each cytokine is secreted and/or sensed by the T cells and is carried between compartments by blood and lymph (Fig. 2a). **At the cellular scale**, CD4+ T cells are described by an agent-based model. In the activation stage, a naive agent migrates from the target organ to the lymphoid tissues (Fig. 2a) where it is stimulated until activation into an effector agent [9–11]. In the expansion stage, it changes its phenotype adaptively in response to changes in cytokine concentrations, undergoes cell cycling, produces cytokines, and migrates to the target organ [8,12,13]. In the contraction stage, it undergoes activation-induced cell death (AICD) [14], activated cell-autonomous death (ACAD) [14], or memory formation, giving rise to a memory agent in the final case [15]. After the contraction stage, an agent representing a memory CD4+ T lymphocyte behaves as a naive agent and goes through the same three stages, albeit with different activation conditions and locations. **At the molecular scale**, signal transduction and gene regulation in each agent are described by a logical model consisting of 73 Boolean variables representing proteins and genes, and 156 pairwise interactions between these variables. This model is an extended version of a previously published model used to study the plasticity of CD4+ T cell differentiation [8]. Based on the generic human metabolic model Recon 2.2.05 [16], together with 159 microarray datasets and 20 proteomic datasets, five phenotype-specific constraint-based models were built. The metabolic networks of effector Th0, Th1, Th2, Th17, and Treg CD4+ T cells comprise 4234, 4160, 4674, 5223, and 3854 reaction fluxes respectively, and 2452, 2377, 2623, 2833 and 2178 metabolites respectively. Cytokine concentrations are used as inputs for the cellular and molecular models; they are conversely influenced by the cellular models. The models at the molecular scale determine most of the attributes of the cellular agents.

**Figure 2:**
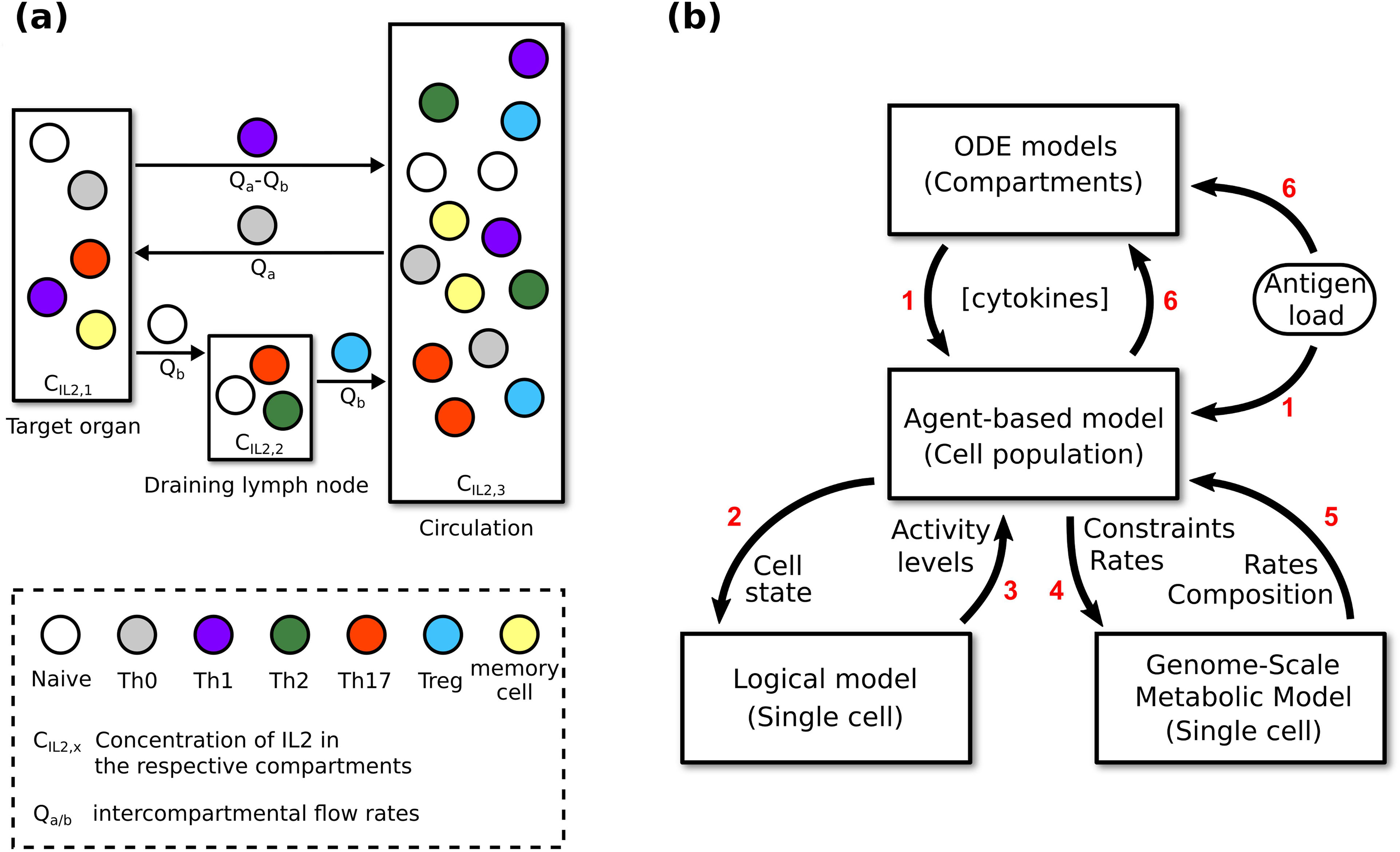
information flow within the multi-scale model. (a) Active migration of agents and passive transport (carried by blood and lymph) of cytokines between compartments representing different organs. See Online Methods for details of how the intercompartmental flow rates affect the cytokine concentrations. The target organ varies depending on the infection being modeled. The cells depicted in each compartment are only for illustrative purposes. See Online Methods for more information on the distribution of cells in each compartment. (b) Flow of information between the models during simulations. Red numbers indicate the sequential steps of a complete update at the systemic/cellular time scale. Labels on the arrows list the variables exchanged between models.

Different parameter sets can be used to model and simulate diverse immune phenomena. At the highest level, the simulation algorithm is a series of fixed time steps, the number and duration of which are determined by the duration and resolution of the modeled event. Fig. 2b illustrates the information flow between the four components of the multi-scale model. Every time step, each agent is evaluated for all behaviors according to the relevant logical model outputs, such as activation (naive agents only), cell death, memory formation, migration, and cytokine secretion. Its progression through the mammalian cell cycle is dependent on various metabolic/synthesis rates. For each agent, the logical model is implemented by using information from the agent and the compartment it is in to parameterize the input components, then calculating the activity level of each non-input component at convergence (estimated by the Kolmogorov-Smirnov test [17]), and finally using the outputs to update the agent’s attributes and inform their behaviors. Some of the outputs are used to parameterize the constraint-based models in order to update the agent’s metabolic/synthesis rates. Cytokine concentrations are then calculated by considering the attributes of every agent. A more detailed account can be found in the Supplementary Text.

### The model reproduced observed responses of naive CD4+ T cells to different combinations of cytokines

Eizenberg-Magar *et al.* [18] used different combinations and dosages of cytokines to differentiate naive CD4+ T cells into effector cells of various phenotypes. IL12 produced the Th1 phenotype, a combination of IL2 and IL4 the Th2 phenotype, a combination of IL6 and TGFβ the Th17 phenotype, and finally, a combination of IL2 and TGFβ resulted in a Treg phenotype. A summary of this study can be found in the Supplementary Text. We validated the multi-scale model by reproducing these results. We conducted four Monte Carlo simulations, each comprising 50 realizations of the modeled system (standard specifications, see Online Methods). In each case, the initial cytokine concentrations were configured to match a set of concentrations used in the experimental study. All the other parameters were set according to the standard specifications. We also conducted a fifth simulation with a combination of all five cytokines plus IFNγ. For each simulation, the 50 realizations were averaged to generate the results presented in Fig. 3a.

**Figure 3:**
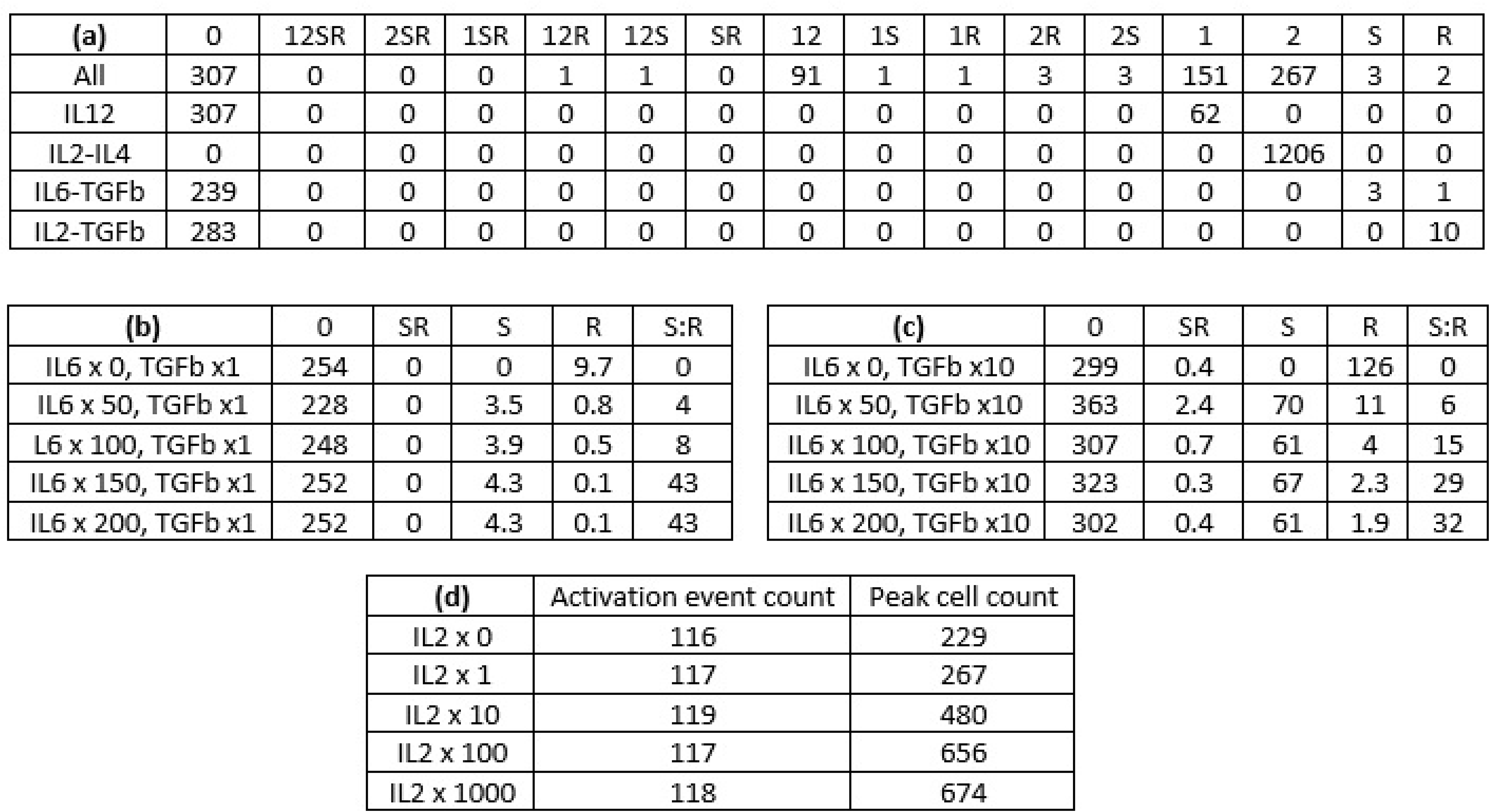
peak CD4+ T cell responses in the presented simulations. Each row represents the peak CD4+ T cell response in a Monte Carlo simulation, where 50 realizations of the modeled system were generated and averaged (see Online Methods and Supplementary Text for details). The peak is defined as the time point with the maximum number of effectors and reactivated memory cells. In the column labels, 0 represents Th0; 1, Th1; 2, Th2; S, Th17; and R, Treg. A combination of symbols represents a mixed phenotype; for example, 12 represents the Th1-Th2 hybrid phenotype. (a) Phenotypic distributions of CD4+ T cells in different cytokine media. The initial cytokine concentrations were set to match an experimental study [18]. The first row represents the case with IL2, IL4, IL6, IL12, IFNγ, and TGFβ. (b) Phenotypic distributions of CD4+ T cells given different initial concentrations of IL6 (multiples of 20 ng/mL). In each simulation, the initial concentration of TGFβ was set to 10 ng/mL, while the other cytokine concentrations were set to zero. S:R denotes the ratio of Th17 count to Treg count (c) Same as (b), except the initial concentration of TGFβ was set to 100 ng/mL. (d) Total counts of naive cell activation and maximum cell counts given different initial concentrations of IL2 (multiples of 5 ng/mL). The initial concentrations of the remaining cytokines were set to zero.

As illustrated in Fig. 3a, the observed differentiation patterns, as summarized in the last paragraph, were reproduced *in silico*. Such differentiation patterns were considered for non-physiological cytokine concentrations at the single-cell level in our previous study [8]. The multi-scale model replicated our previous results for physiologically relevant cytokine concentrations at the cell-population level. The sensitivity of the multi-scale model varies for different cytokine combinations. For example, IL2 and IL4 triggered a strong response, converting all naive CD4+ T cells into a uniform population of Th2 effector cells. In contrast, the Treg response to IL2 and TGFβ was much weaker, with most effector cells adopting the Th0 phenotype. Fig. 4a offers insights into the logical model attractors corresponding to the five combinations of cytokines. It presents the activity levels of selected nodes in estimated attractors, which are consistent with the results in Fig. 3a. For example, *GATA3* (activity level of GATA3) is 1 (maximal) for the IL2-IL4 combination, thus explaining the strong Th2 response presented in Fig. 3a. Details regarding how these estimated attractors were computed can be found in the Supplementary Text.

**Figure 4:**
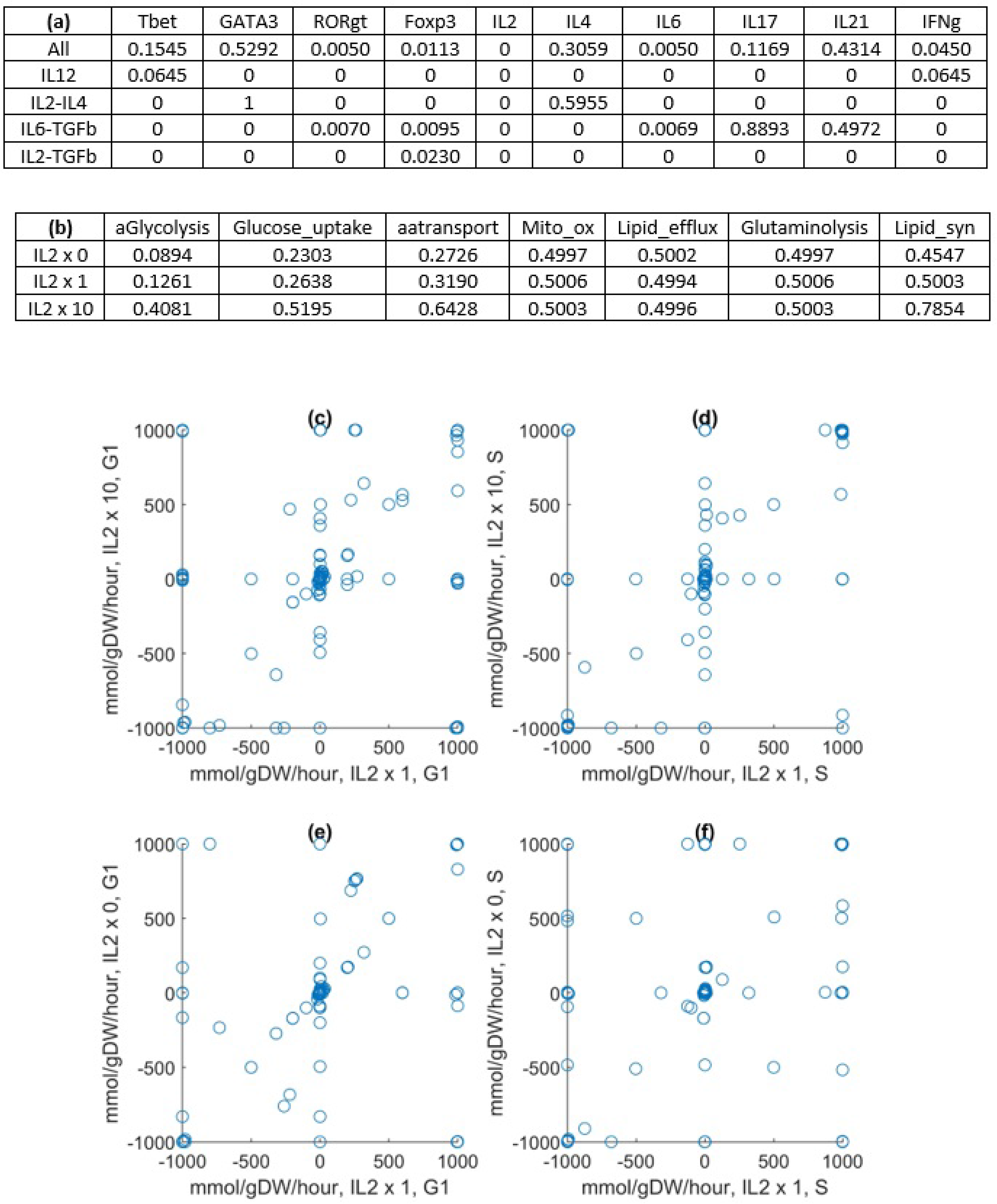
representative results from the models at the molecular scale. Details about how the molecular models were used on their own can be found in Online Methods and Supplementary Text. (a) Activity levels of selected components of the logical model in the estimates of attractors corresponding to different cytokine inputs. (b) Activity levels of metabolic components of the logical model in the estimates of attractors corresponding to different IL2 input levels (0, 5, and 50 ng/mL). (c)-(f) Scatter plots comparing the metabolic fluxes in a Th0 cell during different cell cycle phases (G1 and S) at different IL2 concentrations (0, 5, and 50 ng/mL).

During our simulation corresponding to the IL6-TGFβ combination, both Th17 and Treg phenotypes were adopted by effector cells. Although TGFβ alone induced T cell differentiation into the Treg lineage, when IL6 was also present, T cells differentiated into Th17 or Treg in an IL6 concentration-dependent manner. The simulation with IL6 and TGFβ was repeated with multiples of the original IL6 concentration (20 ng/mL) used by Eizenberg-Magar *et al*. [18]. The results show a positive correlation between the Th17:Treg ratio and IL6 concentration (Fig. 3b), in agreement with our previous dose-response study [8]. Repeating the simulation with a tenfold higher concentration of TGFβ resulted in higher cell counts but the same general trend (Fig. 3c).

### The model reproduced IL2-mediated metabolic regulation of CD4+ T cells

IL2 is a key cytokine in the metabolic regulation of CD4+ T cells because it induces both their proliferation and activation-induced cell death [19]. Therefore, this validation study centered on the regulatory effects of IL2 on the expansion stage of the cell. Monte Carlo simulations were carried out with different initial IL2 concentrations (0, 5, 50, 500 and 5000 ng/mL, multiples of the concentration used in experiments [18]). All other initial cytokine concentrations were set to zero. The other parameters were set according to the standard specifications (Online Methods).

The results (Fig. 3d) show that with no IL2 in the compartments, 116 activation events (out of 433 potential activation events, one for each naive T cell) had occurred when the simulated immune response peaked, but 229 effector cells and reactivated memory cells were present at the peak, suggesting a basal level of proliferation even in the absence of IL2. At the highest IL2 concentration, the maximum number of cells (674) is almost three times that in the absence of IL2 (229). These results can be compared to the experimental results in a study [20] where CD4+ T lymphocytes were stimulated with anti-CD3 and IL2 at various concentrations (0.625 to 25 ng/mL). At the highest IL2 concentration, the extent of DNA replication in the cells undergoing the first round of division (a proxy for cell count) was measured at almost three times that at the lowest IL2 concentration, in agreement with our simulation results.

The population-level results shown in Fig. 3d can be explained by considering the models embedded in each agent at the molecular scale. Fig. 4b shows the activity levels of selected nodes in the estimated attractors corresponding to three initial IL2 concentrations (0, 5, and 50 ng/mL). The results indicate that the activity levels of metabolic nodes should either increase or stay constant in response to higher IL2 concentrations. Details regarding how these estimated attractors were computed are presented in the Supplementary Text. The activity levels of metabolic nodes in each estimated attractor were used to alter the constraints on the metabolic fluxes in the Th0 metabolic model before the fluxes were optimized for both the G1 and S phases of the cell cycle. Mathematically and computationally, it means that an objective function representing the goal of each phase was defined by combining the fluxes, and it was maximized by finding an optimal set of fluxes under the aforementioned constraints (linear programming problem solved by the simplex algorithm [16]). The results are shown in Fig. 4c to Fig. 4f. Each scatter plot shows multiple fluxes off the identity diagonal, indicating changes of flux values, and even fluxes lying on the opposite diagonal, indicating reversals of the corresponding fluxes. Biologically, it means IL2 affects metabolism in a high nonlinear manner, implying drastic and counter-intuitive changes in the growth dynamics of CD4+ T cells. For example, it boosts the biosynthetic fluxes related to aerobic glycolysis and lipid synthesis.

### The model reproduced the population dynamics of CD4+ T cells during influenza infection

We used the standard specifications of the model to simulate the CD4+ T cell response to influenza infection. Due to the high computational costs associated with linear programming and exponential population growth, the initial number of naive agents in the draining lymph node was set to 10 to represent a portion of the node only. The initial cell counts in the other two compartments were set to respect the relative population sizes in the three compartments. There were no more sophisticated reasons behind the choice of 10. As for the influenza infection, the viral trajectory from an experimental study [13] was converted into the input signal (Fig. 5a, the curve labeled 0.8) discussed in the Online Methods and, with more detail, in the Supplementary Text. The signal ranges from 0 to 1 in arbitrary units. Biologically, it represents the viral load and the activity of the immune system triggered by the virus and relevant to CD4+ T cell activation. Computationally, it is the probability of its stimulating a CD4+ T cell agent.

**Figure 5:**
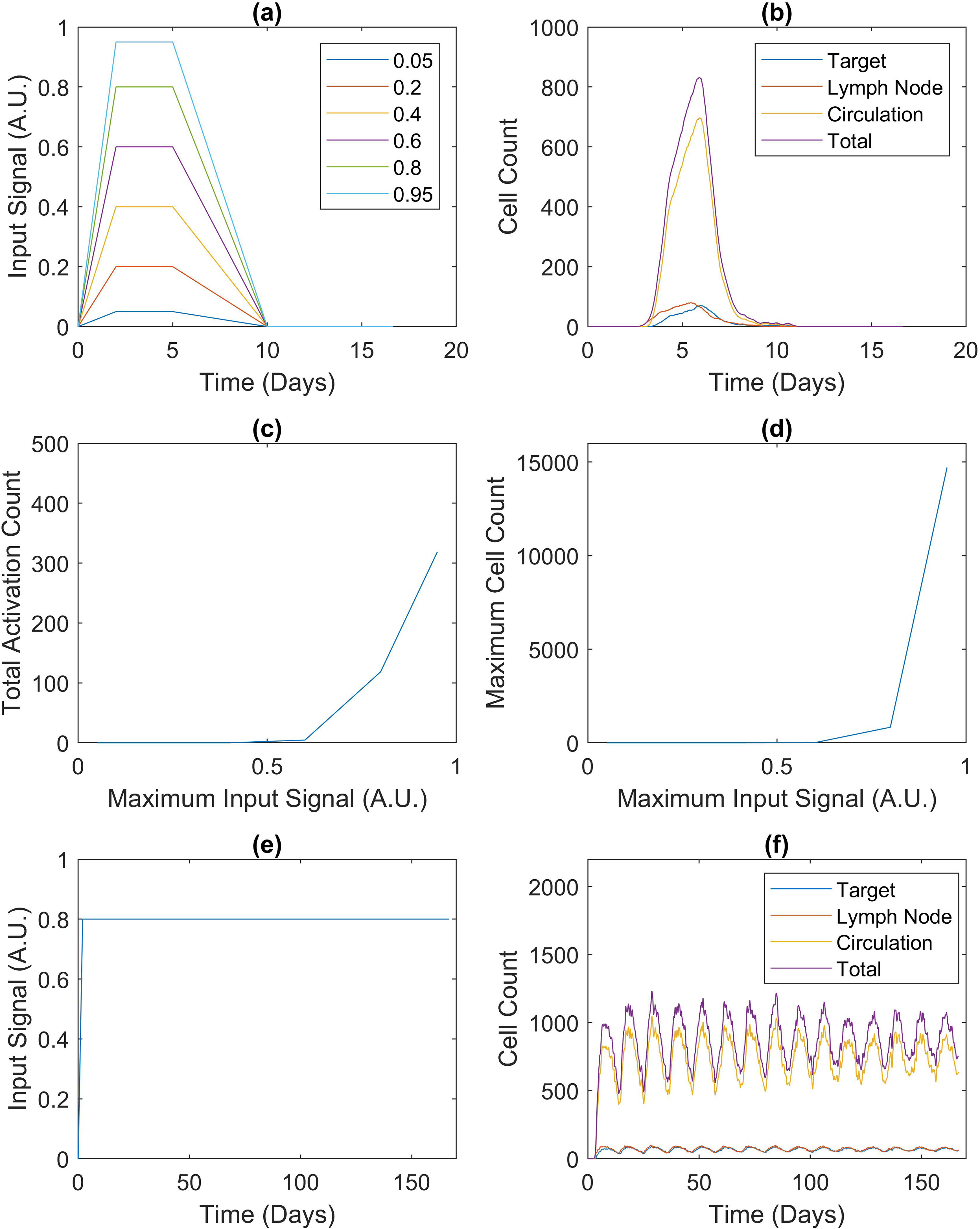
CD4+ T cell responses to different infections. CD4+ T cell responses involve effectors and reactivated memory cells only. (a) Input signals representing acute infections with different peaks. The signal strength is in arbitrary units (A.U.). The signal labeled 0.8 represents the viral trajectory from an experimental study about influenza [13] and is known as the default input signal in this paper. (b) Population dynamics in response to the default input signal, the average of 50 realizations of the modeled system. (c) and (d) Total counts of naive cell activation and maximum cell counts corresponding to the input signals in (a). (e) Input signal representing a hypothetical chronic infection. (f) Population dynamics in response to the input signal in (e). Details about the signals and simulations can be found in Online Methods and Supplementary Text.

The simulated population dynamics are shown in Fig. 5b. In the first four days post-infection, the trajectory representing the draining lymph node rises nearly five-fold (10 to almost 50). In comparison, over the same period, the experimental study reported a 16-fold expansion of the corresponding population in the draining mediastinal lymph node. Although the rise in cell count in Fig. 5b is smaller than the observed expansion, the same qualitative trend is present in both sets of results. The simulations also indicate that the CD4+ T cell population in the lungs should rise more slowly than that in the lymphoid tissues: a third of the peak after four days of infection. The predicted time delay is in qualitative agreement with the experimental study, where no CD4+ T cells were detected in any lung tissues at four days post-infection. In the experimental study, all cell populations were observed to peak at six days post-infection and decline between six and eight days post-infection, again agreeing with Fig. 5b.

### The model predicted switch-like and oscillatory emergent behaviors of CD4+ T cells in response to infection

After validation of the multi-scale model, we performed further simulations to test its robustness in response to different input signals. To this end, the default input signal was modified in terms of its peak strength and long-term behavior as explained in the Supplementary Text. In other words, we changed the signal quantitatively and qualitatively.

The first robustness test centered on the input signal strength. We generated five input signals to model acute infections with different peak strengths (Fig. 5a). The Monte Carlo simulation presented in the previous subsection was repeated with each signal. Fig. 5c and 5d summarize the results of averaging the six sets of 50 realizations. During the simulations, the input signals peaking at 0.05, 0.2, and 0.4 were too weak to activate any naive cells. In the model, a naive cell must migrate to the draining lymph node before activation can occur, but it can only move and stay there in response to a strong input signal, effectively forming a filter for weak input signals. On the other hand, when the input signal was set to peak at or above 0.6, the effector cells amplified its effects on proliferation nonlinearly. In Fig. 5a, as the input signal peak rises from 0.6 to 0.8 to 0.95 (less than two-fold), the number of activation events increases from 4.60 to 118.46 to 318.58 (almost 70-fold), and the maximum cell count increases from 7.70 to 831.84 to 14705 (over 1900-fold). In the model, each effector cell’s activation level is an attribute reflecting the input signal, and it is passed to the nonlinear logical model, which ultimately regulates proliferation. Cooperatively, the effector cells secrete IL2 to boost further proliferation in the population, so as they proliferate, the rate of proliferation increases. This effect is enhanced by the simple fact that each cell produces multiple daughter cells. Taken together, our results suggest that CD4+ T cells demonstrate a switch-like behavior in response to infection.

The second robustness test centered on the input signal’s pattern. To simulate chronic infection, we produced a steady input signal which does not decrease after peaking (Fig. 5e). Unlike the acute input signal (Fig. 5a), which led to a decrease in the simulated immune response (Fig. 5b) after the signal was removed, the chronic input signal produced oscillations in the population size of CD4+ T cells (Fig. 5f). This behavior emerges from the interplay between persistent stimulation and episodic replenishment of naive CD4+ T cells. At the beginning of each cycle, the input signal activates naive cells into effector cells. As more effector cells enter the contraction stage, cell death occurs more and more frequently, while the supply of naive cells continues to drop, slowing down the growth of the effector cell population. Eventually, their death rate exceeds their replacement rate, leading to a decline in their population. When the naive cell population drops below a threshold, a fixed number of naive cells are added to the compartments to initiate a new cycle. Oscillations can also be seen in the concentrations of secreted cytokines (Fig. 6). Comparison with the first row of Fig. 4a explains the variations between the six sets of concentration dynamics. *IL6* is extremely low in the relevant estimated attractor. This, together with the fact that the cytokine-secreting ability of an effector CD4+ T cell increases with its division count [21,22], explains why the target organ concentration is higher than the lymph node concentration in Fig. 6c (more divided cells in the target organ and low secretion rate in the lymph node). In the other panels of Fig. 6, the lymph node concentrations are higher because of the lymph node’s small volume, while the activity levels of the relevant cytokine nodes are not low enough to offset this effect. Fig. 6a seems to contradict *IL2* being zero in the estimated attractor, but the relatively plastic effector cells in the lymph node secrete IL2 by default [21,22]. Finally, the concentrations of IL17 (Fig. 6d) and IL21 (Fig. 6e) are an order of magnitude or two lower than the rest but fluctuate more. It is because their initial concentrations were set to zero.

**Figure 6:**
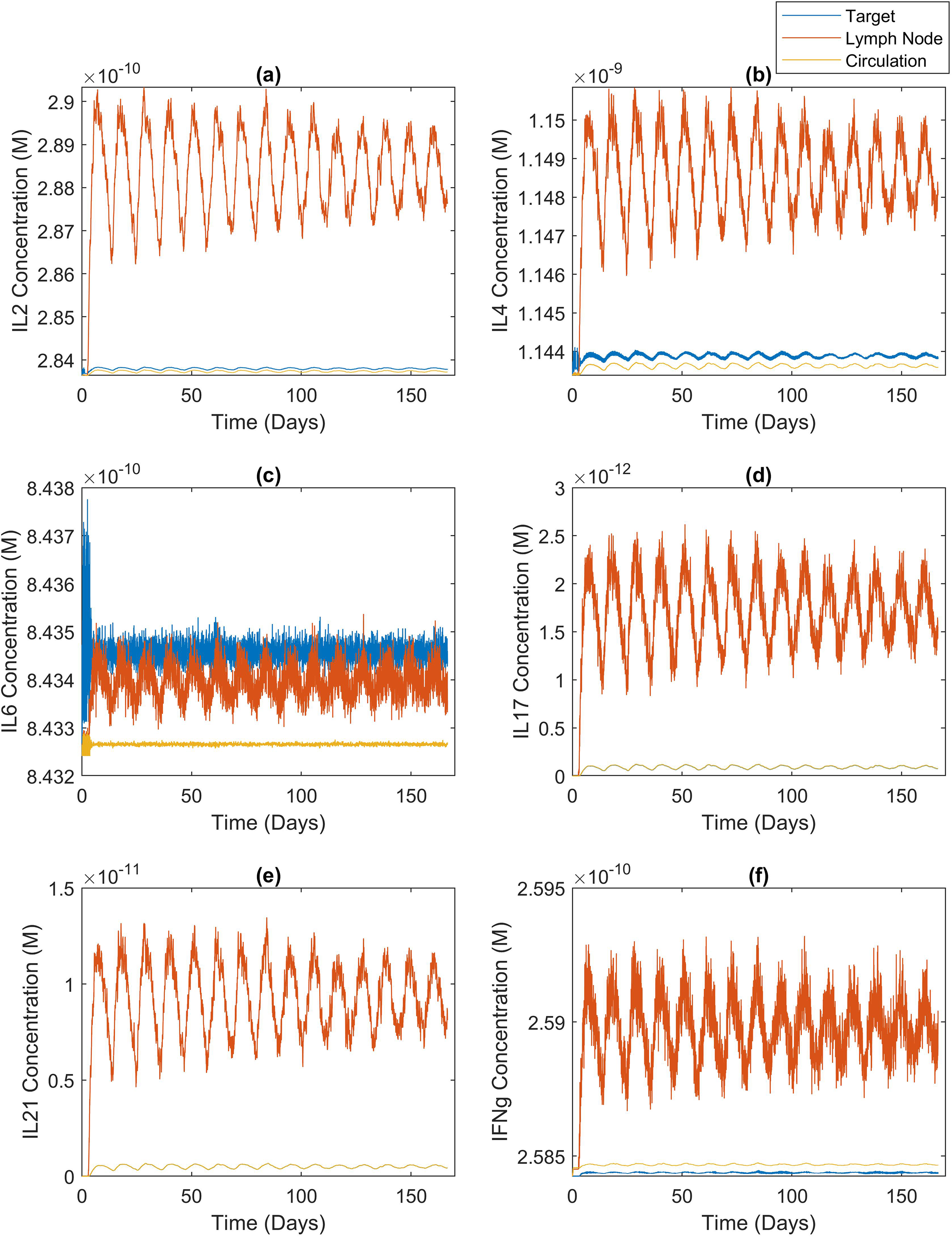
cytokine concentration dynamics during chronic infection. Concentration dynamics of the six cytokines secreted by CD4+ T cells during a hypothetical chronic infection. Details about the signal and simulation can be found in Online Methods and Supplementary Text.

## Discussion

Numerous computational approaches have been used to model the immune system and its functions [23], particularly in relation to T cells. Although impressive and informative, those studies did not cover the scales present in our model, include multiple compartments, or use a suitable modeling approach for each scale. As such, they could not capture essential nonlinear aspects of immune responses, such as emergent properties. Many models are phenotypic in that they lack precise molecular and cellular details, so they cannot readily be informed by-and their outputs compared with-experimental and patient data [24]. Some modeling studies used detailed agents but each focused on a given compartment such as germinal centers [25]. A few agent-based models were built by including different immune cell types in multi-compartment setups not dissimilar to the one presented in this paper [26,27]. However, the agent behaviors were not linked through the use of adaptive, intracellular models of biochemical networks. While they made the individual agents adapt to the changing cell populations and cytokine concentrations, they did not capture the resulting feedback control with the overall models. Logical models informing the phenotype of CD4 T cells have been developed [28] and even been used to build multi-cellular models [29]. However, if used for our study, the lack of differentiated anatomical compartments would not allow time delays caused by cell migration at the systemic scale to be captured, such as the time delay in Fig. 5b. As such, they would not display the switch-like behavior which depends on the fact that CD4+ T cells must migrate to the draining lymph node before being activated. Finally, while metabolism has been shown to be essential for T cell activation [30] and genome-scale models of CD4+ T cells exist [6], they were not used in any dynamical multi-scale modeling studies.

To the best of our knowledge, our model — through the use of four different but integrated modeling approaches — is the first using a compartmental approach to understand immune responses at three different spatial scales. In addition to tackling the drawbacks outlined above, the hybrid and multi-scale approach confers the advantage of modularity on the model. New agent types can be developed to represent other cell types (such as B cells), and integrated with the existing framework. This was how we expanded our existing logical model of CD4+ T cell differentiation [8], and integrated it with the framework to reproduce simulation results (at the single-cell level and for abstract cytokine levels) at the population level using physiological cytokine concentrations. Modularity also permits reusability of existing models; new versions of intracellular models, using similar or different modeling approaches, can be integrated with the framework with minimal effort. In addition to the gains in computational performance brought by using different levels of granularity for different parts of the model, one could decouple the parts and cache simulation results. For example, we built a library of steady states for our metabolic models to match different estimated attractors of the logical model. Finally, modularity also helps with data integration. For instance, if a dataset related to memory formation is available at the cellular level, while another dataset about metabolism is available at the molecular level, one can easily work with both datasets within our framework.

The multi-scale nature of our model is very important because immune responses are multi-scale. The concentrations of cytokines at the systemic scale, the collective behavior of the cell population, and the heterogeneity within the population are interwoven due to nonlinear structures such as feedback loops. For example, the difference between our results in Fig. 3d and an experimental study [20] seems to indicate that the cells in the experiment were more sensitive to IL2 than the agents in our model. However, the effects of activation-induced cell death reported in [31] were not measured in the experiment which only considered the first round of division. In contrast, our model captures the complex relationship between IL2 (systemic scale), cell expansion (cellular scale), and AICD (molecular scale): IL2 stimulates both cell expansion and death. The dampening effects of AICD on cell expansion can be seen in the saturating behavior illustrated by Fig. 3d.

At the systemic level, compartmental ordinary differential equations were chosen to model cytokine dynamics since the compartments are considered isotropic and molecules are indistinguishable. Because of the very large number of molecules, they can be solved using deterministic algorithms. At the cellular level, an agent-based approach was selected to serve the needs to model heterogeneity in the population and track each cell’s phenotype. It was decided that cell expansion should be treated as a process (mammalian cell cycle) rather than a single event (division) to provide realistic links between the agent-based model and the models at the molecular level. In our simulations, for most agent behaviors (such as differentiation), the activity levels of the logical model were normalized before being passed to the agent-based models as probabilities. On the contrary, the activities of the nodes interfacing with the genome-scale metabolic models were used to alter the constraints on metabolic fluxes before the fluxes were optimized. A constraint-based approach was chosen over ordinary differential equations for the metabolic models to circumvent the lack of kinetic parameters and to allow a comprehensive (genome-scale) representation of metabolism. A disadvantage is the lack of transient dynamics. However, our algorithm mitigates the problem by updating the steady-state for each cell at every time step. Because this solution comes with a high computational cost, we built a library of optimized fluxes corresponding to the estimates of common logical model attractors.

Our model was able to reproduce diverse experimental observations made on CD4+ T cells, so we are confident in the novel but plausible properties it predicted. The model reproduced the ability of CD4+ T cells to differentiate into different phenotypes in a context-dependent manner, as well as the regulation of CD4+ T cell metabolism by IL2. We also reproduced the population dynamics of CD4+ T cells during influenza infection. These experimental observations were made in completely independent studies, and we emphasize that, except for the cytokine concentrations, they were not used to parameterize the model. The emergent properties we identified in virtual CD4+ T cell populations, switch-like and oscillatory behaviors, both arise from interactions between biological phenomena taking place at different spatial scales. The switch-like behavior might help the body prevent chronic inflammation in the continuous presence of a small and non-threatening amount of antigen. In fact, switch-like behaviors have been observed in populations comprising other immune cell types. For instance, the chronic inflammation triggered by mast cells is a bistable switch [32]. Similarly, oscillations are not uncommon in the immune system [33], possibly as side effects of homeostasis. However, no one has observed or predicted such nonlinear behaviors in relation to CD4+ T cells. This fact highlights our model’s ability to capture the complexities of the immune system.

We plan to add other immune cell types to the framework, as well as a method to model physical cell-cell interactions. This should provide a virtual immune system, well-suited to exploring the fundamental properties of the immune system and to modeling immune responses to specific diseases. Just as we found switch-like and oscillatory behaviors of CD4+ T cells, its multi-scale nature will allow researchers to unravel counterintuitive, nonlinear properties in the broader immune system.

## Online Methods

### Structure of the multi-scale model

#### Systemic scale

The multi-scale model is divided into three isotropic compartments. In each compartment, IL2, IL4, IL6, IL12, IL17, IL18, IL21, IL23, IL27, IFNγ, and TGFβ are represented as time-dependent homogeneous concentrations. An agent representing a CD4+ T cell can sense all cytokines except IL17 and IL21, and it can secrete IFNγ, IL2, IL4, IL6, IL17, and IL21. The concentrations of the former are passed to the models at the cellular and molecular scales (see below), while the concentrations of the latter are influenced by those models. This setup is consistent with our previous study at the molecular scale [8]. Furthermore, in each compartment, a study-dependent and time-varying input signal represents the antigen load and the activity of the rest of the immune system. Cytokine concentrations are governed by 33 ordinary differential equations:

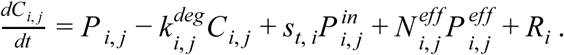

*C*_*i,j*_ is the concentration of cytokine j in compartment i (M), *P*_*i,j*_ is the baseline production rate (M·h^−1^), 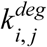 is the degradation rate constant (h^−1^), *s*_*t, i*_ is the input signal (antigen load and the rest of the immune system, dimensionless) in the compartment at time t (hours), 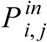 is the production rate in response to the input signal (M·h^−1^), 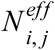 is the number of cells/agents producing the cytokine in the compartment, 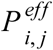 is the rate of production by one cell/agent in the compartment (M·h^−1^), and *R*_*i*_ is a transport term (M·^h−1^). Compartment 1 is the target organ, 2 the lymphoid tissues, and 3 the circulatory system. The transport terms are described by the following equations:

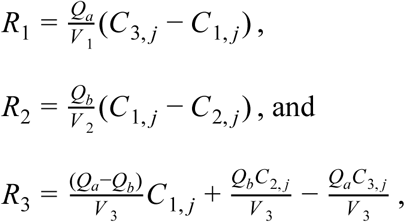

where *V* _1_, *V* _2_, and *V* _3_ are the compartments’ volumes in dm^3^, *Q*_*a*_ is the rate of blood flow into the target organ (dm^3^·h^−1^), and *Q*_*b*_ is the rate of lymph flow into the circulatory system (dm^3^·h^−1^).

#### Cellular scale

Cells are represented as discrete and autonomous agents defined by a set of attributes and behaviors. These agents respond to changes in both the cytokine concentrations and the input signal. A newly created agent represents a naive CD4+ T cell. They are created in all three compartments. Then, they undergo three major life cycle stages: activation, expansion, and contraction.

During the <u>activation stage</u>, a naive agent exhibits three behaviors. First, in response to the input signal in the lymphoid tissues (*s*_*t*, 2_), the naive agent migrates from the target organ to the lymphoid tissues at a higher rate and stays there. Second, *s*_*t, 2*_ is the probability that the naive agent is stimulated in one time step, and if it gets stimulated, its *TCR_stimuli* increases by one, and it records the signal magnitude (between zero and one) and cytokine concentrations. Third, if the stimulation is adequate (*TCR*_*stimuli* = *act*_*thres*), it is activated and becomes an effector agent. Computationally, activation involves using the logical model and appropriate metabolic models to determine the effector agent’s attributes, such as its phenotype (Th0, Th1, Th2, Th17, Treg, and hybrid phenotypes comprising these five states).

During the <u>expansion stage</u>, an effector agent exhibits four behaviors. First, it continuously monitors the environment for changes in the input signal and cytokine concentrations, and updates internal attributes, such as the amount of biomass, which are used to track its progress through the cell cycle. Second, at the end of the cell cycle, it divides into two identical copies. Third, it secretes cytokines in accordance with its changing cellular phenotype. Fourth, after several rounds of division, it leaves the lymphoid tissues, enters the circulatory system, migrates unidirectionally to the target organ, and stays there.

During the <u>contraction stage</u>, an effector agent exhibits four behaviors. First, if it is re-stimulated by the input signal, it may undergo activation-induced cell death (AICD). Second, in the absence of restimulation, it may become a memory agent. Third, in the absence of restimulation and memory formation, it may undergo activated cell-autonomous death (ACAD). Fourth, if none of the first three cases applies, its activation level (in terms of *TCR_strength* and *CD28_strength*) will decrease by an amount determined by the input signal’s strength in the time step.

After the contraction stage, a memory agent undergoes the same three stages described above for naive agents. However, fewer stimuli are required for its activation and the resulting activation level is higher [34,35]. Unlike a naive agent, a memory agent can get stimulated and activated in all three compartments. Upon activation, it becomes a reactivated memory agent and behaves exactly like an effector agent [36]. Unless otherwise stated, all properties of effector cells and effector agents apply to their reactivated counterparts too.

#### Molecular scale

Most of the attributes of each agent are determined by a logical model of signal transduction and gene regulation, and constraint-based models of metabolism.

**Signal transduction pathways and gene regulation processes** captured in the model include the canonical pathways regulating the differentiation of CD4+ T cells into various cell phenotypes [8], and events such as aerobic glycolysis, memory formation, and cell death. These pathways are detailed in the Supplementary Text. The network diagram of the logical model is presented in Fig. S1. Its structure is detailed in the Supplementary Text. The model itself is accessible in Cell Collective [37], where it can be downloaded as an SBML file, or directly modified and implemented in simulations.

The logical model has three classes of inputs corresponding to 1) cytokine concentrations, provided by the ordinary differential equations, 2) receptor signaling (*TCR* and *CD28*), determined by the agent’s activation level, and 3) agent attributes transformed into logical model inputs, such as the activity of mTORC1, concentration ratio of AMP and ATP, abundance of ribosome, and the agent’s resistance to apoptosis.

The outputs of the logical model affect agent attributes and behaviors, such as phenotypes (Th0, Th1, Th2, Th17, and Treg), apoptosis, and cytokine secretion. They also parameterize the metabolic models. There are four classes of outputs representing 1) the transcription factors Tbet, GATA3, RORgt, and Foxp3, 2) the cytokines produced by the agent, 3) seven metabolic components providing the interface with the genome-scale, constraint-based models of metabolism, and 4) six cellular events (cell cycle progression, autophagy, memory formation, ACAD, AICD via the Fas pathway, and AICD via the B-cell lymphoma 2 (BCL2) pathway).

**The metabolic networks** of effector Th0, Th1, Th2, Th17, and Treg CD4+ T cells are formalized in five phenotype-specific constraint-based models. 159 microarray datasets were integrated with 20 proteomic datasets to define the gene activity profiles (matrices) of the five phenotypes. The profiles and the generic human metabolic model Recon 2.2.05 [16] were used to build the constraint-based models by the GIMME method in the COBRA toolbox [38]. In summary, the Th0 model contains 4234 metabolic fluxes and 2452 metabolites; Th1, 4160 and 2377; Th2, 4674 and 2623; Th17, 5223 and 2833; and Treg, 3854 and 2178.

During a simulation, at each time step at the cellular and systemic scales, multiple simulations at the molecular scale (logical and constraint-based models) are run. At each discrete update of the logical model, the activation probability of each input component serves as an input to the logical model. Some of the outputs (seven metabolic components) are used to set the constraints in the metabolic models, which are then used to calculate the metabolic/synthesis rates in the agents. A detailed account of the simulation algorithm and the corresponding pseudocode can be found in the Supplementary Text.

### Variables and parameters of the multi-scale model

The multi-scale model has four classes of variables and parameters. They are agent attributes, variables/parameters specific to the compartments, settings for the numerical methods, and four parameters that can be varied for model calibration. All the variables and parameters are presented in Table S1 to Table S10. A detailed description of all four classes and the parametric estimation process is in the Supplementary Text, where it is also explained how the model can be used to study diverse immune phenomena by changing the parameters.

### Convergence study

Because the model contains stochastic elements, such as the logical model and most agent behaviors, multiple simulations (realizations of the model) must be performed to obtain robust averages of model outputs. A convergence study helps the modeler decide how many simulations are required to guarantee robustness. It is necessary following parameterization to describe a specific immune phenomenon. The convergence study we ran before the validation and robustness studies presented in this paper, as carried out using the default, acute-infection-like, input signal (Fig. 5a, curve 0.8) is described in the Supplementary Text. Based on the results, we decided that 50 simulations were required for the average model outputs to converge. The specifications detailed in the supplementary Text, namely the input signal, initial cytokine concentrations, cytokine production rates associated with the input signal, and the need for 50 model realizations, are the standard specifications of the studies presented in this paper. In Results, it is understood that a simulation refers to a set of 50 realizations of a parameterized model.

### Source code

The model source code is freely available on GitHub (https://github.com/HelikarLab/multiscale-CD4Tcells).

## Acknowledgements

This work was supported by NIH grant 1R35GM119770-04 to T.H. and by the Systems Biology Grant of the University of Surrey to M.B.

## Author Contributions

T.H. conceived the study. K.Y.W., M.B., and T.H designed the study. K.Y.W. built and implemented the model, performed the validation studies, and with the help of M.B., carried out the robustness studies. B.L.P., with the help of A.R.S., categorized the fluxes in the metabolic models into groups matching the metabolic nodes in the logical model. A.L.F. helped K.Y.W. design the compartmental setup. K.Y.W., M.B., and T.H. analyzed the data and then hypothesized about the presence of switch-like and oscillatory behaviors. K.Y.W., B.L.P., M.B., and T.H. wrote the manuscript. M.B. and T.H. provided scientific leadership by supervising the study.

