## Supplemental Information for "Multi-Approach and Multi-Scale Model of CD4+ T Cells Predicts Switch-Like and Oscillatory Emergent Behaviors in Inflammatory Response to Infection"

### Supplementary Information

#### *Supplementary text*

*1. Logical model of signaling pathways and gene regulation.*
*2. Signaling pathways that regulate metabolism.*
*3. Signaling pathways that regulate apoptosis and memory formation.*
*4. Summary of variables and parameters for the multi-scale model.*
*5. Simulation algorithm.*
*6. MATLAB program used to implement the multi-scale model in order to simulate the evolution of agents and cytokine concentrations in response to an input signal.*
*7. Default input signal.*
*8. Input signals representing hypothetical infections.*
*9. Details on the cytokine combinations and dosages used in the presented simulations.*
*10. Convergence study.*
*11. Estimating logical model attractors corresponding to different cytokine combinations and dosages.*
*12. Re-using the multi-scale model to study diverse immune phenomena by changing the parameters, running Monte Carlo simulations, and analyzing the distribution of results.*

#### *Figure S1 - S3*

#### *Table S1 - S10*

#### Supplementary text

##### 1. Logical model of signaling pathways and gene regulation.

The logical model has three classes of input components and therefore three classes of inputs (activation probabilities). During a simulation, the activation probability of each input component is the probability that it (*e.g.*, a cytokine component) is active at the latest time step; the probability is determined by models at the cellular and systemic scales (agent-based model and ordinary differential equations). When we use an italicized name (*e.g.*, *IL2*), we refer to the activation probability of an input component, whose name is normal (*e.g.*, IL2). The first class of inputs contains the activation probabilities of the cytokine components, including *IL2_e*, *IL4_e*, *IL6_e*, *IL12*, *IL18*, *IL23*, *IL27*, *IFNg_e*, and *TGFβ*. The cytokine concentrations are provided by the ordinary differential equations, and the concentrations are compared to the abundance of cytokine receptors in order to determine the activation probabilities. The second input class is concerned with receptor signaling (*TCR* and *CD28*), which is determined by the agent's activation level. T-cell receptor (TCR) and CD28 receptor stimuli are converted into the agent's activation level (an attribute) and then the two logical model inputs. With respect to the third class, other agent attributes are transformed into logical model inputs too; *mTORC1_t* reflects how persistently active the mammalian target of rapamycin complex 1 (mTORC1) is in the agent; *AMP* captures the concentration ratio of adenosine monophosphate (AMP) to adenosine triphosphate (ATP) in the agent; *Ribosome* reflects the abundance of ribosome in the agent; finally, *Resist* indicates how resistant the agent is to apoptosis. During a simulation, at each time step at the cellular and systemic scales, multiple simulations at the molecular scale (logical and constraint-based models) are run. At each discrete update of the logical model, the activation probability of each input component serves as an input to the logical model [Helikar *et al.* 2008, Puniya *et al.* 2018].

An output of the logical model is the activity level of the output component it represents. In fact, the output of every non-input (including output) component is its activity level: the fraction of discrete updates when the component is active [Helikar *et al.* 2008, 2012, Helikar and Rogers 2009, Puniya *et al.* 2018]. We refer to the activity level of a non-input component by an italicized name (*e.g.*, *Pro_GATA3*) and the component itself by a normal name (*e.g.*, GATA3). The logical model outputs inform agent attributes (cell fates) and behaviors (cell decisions), such as phenotypes (Th0, Th1, Th2, Th17, and Treg), apoptosis, and cytokine secretion, and/or parameterize the metabolic models as described in a later section, so they serve as inputs to other parts of the multi-scale model. There are four classes of output components and hence four classes of outputs. In the first class, four output components represent the transcription factors Tbet, GATA3, RORgt, and Foxp3, which characterize the phenotypes Th1, Th2, Th17, and Treg, respectively [Puniya *et al.* 2018]. If a cell's phenotype is Th0, it does not mean that its transcription factors are all inactive. For example, consider an agent whose *Pro_Tbet* is 0; *Pro_GATA3*, 0.5; *Pro_RORgt*, 0; and *Pro_Foxp3*, 0. There is a 50 % chance that it represents a cell with the Th2 phenotype and a 50 % chance for the Th0 phenotype. In the second class, six output components represent the cytokines produced by the agent: IL2, IL4, IL6, IL17, IL21, and IFNg [Puniya *et al.* 2018]. In the third class, there are seven metabolic components: aGlycolysis represents aerobic glycolysis in the agent; Glucose_uptake, glucose uptake from its environment; aatransport, amino acid transport between the agent and its environment; Mito_ox, various mitochondrial oxidation events in the agent; Lipid_efflux, secretion of lipid into its environment; Glutaminolysis, catalyzed disintegration of glutamine in the agent; and Lipid_syn, lipid synthesis in it. They provide an interface with the constraint-based, genome-scale metabolic models. In the fourth class, six components represent cellular events: Cycle represents cell cycle progression; Autophagy; Mem, memory formation; ACAD, activated cell autonomous death; AICD1, activation-induced cell death (AICD) via the Fas pathway; AICD2, AICD via the B-cell lymphoma 2 (BCL2) pathway.

#### 2. Signaling pathways that regulate metabolism.

TCR signaling alone can stimulate a myriad of downstream species that regulate an effector cell's metabolic profile, including cMyc, the estrogen-related receptor alpha (ERRa), liver X receptors (LXRs), and the sterol regulatory element-binding protein 2 (SREBP2) [MacIver *et al.* 2013].

The PI3K/AKT/mTOR pathway is important to the effector cell's metabolic regulation [MacIver *et al.* 2013]. PI3K stands for phosphatidylinositol-4,5-bisphosphate 3-kinase; AKT, protein kinase B; mTOR, mammalian target of rapamycin. PI3K is activated by three independent pathways. First, CD28 and TCR cooperatively switch it on [Shi and Sun 2015]. Second, IL2 and the nuclear factor of activated T-cells (NFAT) cooperatively switch it on [Benczik and Gaffen 2004, Rudensky *et al.* 2006]. Third, IL2, IL4R, and NFAT cooperatively switch it on [Brenner *et al.* 2008]. In the expanded logical model, IL2 is represented by the IL2_e component rather than the IL2 component because the new interactions are not coupled to those present in the original model [Puniya *et al.* 2018]. Once activated, PI3K activates mTORC2 in the presence of ribosome. PI3K and mTORC2 cooperatively activate AKT, which activates mTORC1.

The AMPK/mTORC1/Ulk1/2 pathway acts in opposition to the PI3K/AKT/mTOR pathway [MacIver *et al.* 2013]. AMPK stands for AMP-activated protein kinase; Ulk1/2, unc-51-like autophagy-activating kinase 1 and 2. There are two independent ways of activating AMPK. First, an increase in the abundance of AMP relative to that of ATP will activate AMPK [Alers *et al.* 2012]. Second, mTORC1 can activate AMPK following prolonged activation of the former. AMPK activates Ulk1/2, which in turn inhibits AMPK. AMPK and Ulk1/2 both inhibit mTORC1, but mTORC1 feedbacks to inhibit Ulk1/2 only.

Working together, TCR signaling, the PI3K/AKT/mTOR pathway, and the AMPK/mTORC1/Ulk1/2 pathway control the metabolic events depicted in the expanded logical model [MacIver *et al.* 2013]. cMyc activates the cycle genes [Wang *et al.* 2011] and Ulk1/2 activates autophagy [Alers *et al.* 2012].

#### 3. Signaling pathways that regulate apoptosis and memory formation.

A newly activated effector cell is resistant to apoptosis [Brenner *et al.* 2008]. Its resistance and TCR signaling cooperatively induce HPK1 catalytic activity [Liu *et al.* 2000, Brenner *et al.* 2005], which suppresses p73 [Lissy *et al.* 2000, Brenner *et al.* 2005, 2008] and reactive oxygen species (ROS) [Pham *et al.* 2004, Brenner *et al.* 2005, 2008].

**BCL2-mediated apoptotic pathway.** After several rounds of division, the effector cell's resistance will go down [Brenner *et al.* 2008]. TCR signaling is then free to activate ROS [Gülow *et al.* 2005, Brenner *et al.* 2008], which suppresses the AKT-induced activation of BCL2 [Hildeman *et al.* 2003, Benczik and Gaffen 2004, Brenner *et al.* 2008]. BCL2 promotes memory formation [Grayson *et al.* 2000, Brenner *et al.* 2008], inhibits ACAD [Hildeman *et al.* 2002, Brenner *et al.* 2008], and inhibits AICD via the BCL2 pathway

[Brenner *et al.* 2007, 2008]. When BCL2 is inactive, ACAD is allowed, and if p73 is active too, AICD via the BCL2 pathway is also allowed [Lissy *et al.* 2000, Brenner *et al.* 2008].

**Fas-mediated apoptotic pathway.** ROS [Gülow *et al.* 2005, Brenner *et al.* 2008], TCR signaling [Refaeli *et al.* 1998, Brenner *et al.* 2008], NFAT [Rudensky *et al.* 2006], and IL2 [Refaeli *et al.* 1998, Hwang *et al.* 2005, Brenner *et al.* 2008] cooperatively activate the Fas signaling pathway; this process is enhanced when IL4R is active [Brenner *et al.* 2008]. After that, if p73 [Lissy *et al.* 2000, Brenner *et al.* 2008] and STAT1 [Refaeli *et al.* 2002, Brenner *et al.* 2008] are active too, Fas signaling will trigger AICD independently of the BCL2 pathway [Brenner *et al.* 2008]. When the effector cell is resistant to apoptosis, however, the cellular FLICE-like inhibitory protein (CFLIP) inhibits the Fas signaling pathway [Brenner *et al.* 2008]. When this resistance decreases after several rounds of division [Schmitz *et al.* 2003, Fas *et al.* 2006, Brenner *et al.* 2008], IL2 [Refaeli *et al.* 1998, Brenner *et al.* 2008] and NFAT [Rudensky *et al.* 2006] will suppress CFLIP, thus allowing Fas signaling to switch on and trigger AICD as described above; this way of suppressing CFLIP is enhanced by IL4R [Brenner *et al.* 2008].

#### 4. Summary of variables and parameters for the multi-scale model.

A. The agent-related variables and parameters can be divided into three subclasses: metabolism, signal transduction and gene regulation, and cellular behaviors. The parameters in the first two subclasses are all fixed, but some of the parameters in the last subclass must be defined in a study-dependent manner.
    a. Regarding the first subclass, Table S1 lists the attributes about the metabolic state of a CD4+ T cell and Table S2 lists the parameters that control the behaviors related to these attributes. At each time step, for each effector agent, the synthesis rates are updated and the rates are used to update its composition, which is in turn used to determine which cycle phase the agent is in. The parameters in Table S2 are fixed, so nothing in this subclass needs to be altered in order to set up the model for a specific immune phenomenon.
    b. Regarding the second subclass, Table S3 lists the attributes used to determine the logical model inputs, as well as those determined by the logical model outputs; Table S4 lists the parameters used to turn the former attributes into logical model inputs. *TCR_strength* and *CD28_strength* represent the activation level of an agent. When a naive agent just activates to form an effector agent, they are determined by the strength of stimulation ($s_{t,i}$) throughout the activation stage. After activation, in the absence of restimulation, both attributes drop. They are normalized by *TCR_max* and *CD28_max* before being fed to the logical model. For a newly reactivated memory agent, *TCR_strength* and *CD28_strength* are set to *TCR_max* and *CD28_max* respectively. The attribute *i_sum* represents the abundance of cytokine *i* throughout the activation stage. When a naive agent just activates to

form an effector agent, the nine *i_sum* values are divided by the number of stimuli it has received (*TCR_stimuli*) and the average concentrations are fed to the logical model. In the time steps after that, the compartmental concentrations at the latest time step are fed to the logical model directly. In either case, each cytokine concentration is compared to the receptor count (stochastically between *rep_min* and *rep_max*) to calculate the number of cytokine-receptor complexes. This number is divided by *sat_count* to parameterize the activation probability of the corresponding cytokine input component. *Resist* is inversely proportional to the number of division rounds an agent has undergone (*Divisions*); when it has reached its division limit (*div_lim*), *Resist* is set to zero. *Ribosome* represents the abundance of ribosome in an agent, the proxy for which is the ratio of its protein synthesis rate to the maximum possible rate. Every time after the logical model is implemented, *BN_count* is incremented by one and the resulting *Pro_mTORC1* is used to decide whether mTORC1 is expressed. If it is, *mTORC1_t_count* is incremented by one too. When an agent first activates, the corresponding input (*mTORC1_t*) is zero. When *BN_count* is greater than or equal to *BN_count_thres*, *mTORC1_t_count* is divided by *BN_count* and the result is used to parameterize *mTORC1_t*. All of the parameters in Table S4 are fixed, so this subclass does not require user attention when the model is set up for a specific study.

c. Finally, regarding the third subclass, Table S5 lists the remaining agent attributes, which are only relevant within the agent-based model and do not directly influence the other constituent models of the multi-scale model. Table S6 lists the parameters needed to create an agent, as well as the parameters about the agent population. *Phenotype* is updated according to *Pro_Tbet*, *Pro_GATA3*, *Pro_RORgt*, and *Pro_Foxp3* every time after the logical model is implemented. *Phase* is the cell cycle phase an agent is in and depends on the agent's composition. When a naive or memory agent gets stimulated by the input signal, its *TCR_stimuli* is incremented by one. *Plasticity* is the probability that, after the logical model is implemented, an agent will adopt the outcome to update its *Pro_i* values. When a naive agent first activates, its *Plasticity* is one, but *Plasticity* drops after each round of division. After *mig_thres* rounds, *Plasticity* becomes zero [Brown *et al.* 2004]. At every time step, every agent has a chance of migrating to a different compartment; its migration pattern is given by *k_tot*, *k_toln*, and *k_toc*. The initial number of naive agents in a compartment depends on the study under consideration. If the study involves an influenza infection, the target organ will probably be the lungs and the lymphoid tissues will probably be the closest draining lymph node. Naive CD4+ T cells are replenished in a lifelong process [Parkin and Cohen 2001], so new naive agents are added to the compartments when their total count drops below a threshold (*Refill_thres*). The activation thresholds (*act_thres_naive* and *act_thres_mem*) depend on the size of a time step (*step_size*). If a naive cell requires 10 hours of incubation with an antigen prior to activation, that translates to 10 stimuli for an agent if each time step lasts an hour. If it lasts two hours, however, each stimulus will be worth twice as much, so only five stimuli will be needed for activation. $P_{ex}$ is the final agent parameter that requires explanation. For simplicity, an effector agent produces IL2, IL4, IL6, IL17, IL21, and IFNg in a binary fashion (on at $P_{ex}$ or off). However, the compartments have different sizes, so $P_{ex}$ translates to three values of $P_{i,j}^{eff}$ (M $hour^{-1}$). These three values hold for all six cytokines. In summary, only the numbers of cells in the target organ and lymphoid tissues, as well as the corresponding agent numbers in the compartments, must be determined to set up the multi-scale model for a specific study. Otherwise, the other parameters in Table S6 are fixed.

B. The compartment-related variables and parameters are shown in Table S7 and Table S8. They can be categorized into three subclasses: input signal, chemical kinetics, and sizes.

   a. The first subclass has one member only, the input signal ($s_{t,i}$). It is study-dependent and roughly corresponds to the antigen load in compartment *i* at time *t*. More accurately, it represents the antigen of interest and the associated inflammatory reactions that are not mediated by CD4+ T cells. It increases with the antigen load continuously from zero to one. By default, TCR and CD28 signaling strengths ($TCR_{t,i}$ and $CD28_{t,i}$) are the same as the input signal, but the relations may be changed to tailor the multi-scale model for different studies. For a naive or reactivated effector agent, its value is the probability that it will get stimulated at time *t*.

   b. The second subclass consists of both fixed and study-dependent members. For simplicity, a generic degradation rate constant is used for $k_{i,j}^{deg}$. The initial cytokine concentrations ($C_{i,j,0}$) and the cytokine production rates associated with the input signal ($P_{i,j}^{in}$) are study-dependent, however. The $P_{i,j}^{in}$ values link the input signal to the ODEs about cytokine concentrations. The concentration scale ($C_{j,s}$) is fixed and reflects the cytokine concentration range in the human body.

   c. The third subclass consists of study-dependent members. The compartment volumes are represented by $V_1$, $V_2$, and $V_3$ in $dm^3$, where compartment 1 is the target organ, 2 the lymphoid tissues, and 3 the circulatory system. $Q_a$ ($dm^3$ $hour^{-1}$) is the rate of blood flow into the target organ and $Q_b$ ($dm^3$ $hour^{-1}$) is the rate of lymph flow into the circulatory system. The initial numbers of naive CD4+ T cell agents in the compartments ($N_1$, $N_2$, and $N_3$) and the physiological cell numbers ($N_{1,p}$, $N_{2,p}$, and $N_{3,p}$) must also be decided in a study-dependent manner.

C. Certain parameters are needed for the numerical methods used to implement the multi-scale model. They are listed in Table S9. The number of time steps (*steps*) depends on the time frame of a study, while the time resolution necessary for the study determines the size of each step in hours (*step_size*). Because the multi-scale model is stochastic, it must be implemented multiple times to obtain a distribution of results; *rounds* is the number of times needed for convergence, a study-dependent parameter. The ordinary differential equations about the cytokine concentrations are handled by the ode45 solver (Runge-Kutta method) in MATLAB R2017a; *dis_steps* is a parameter the solver needs. The equations are linear and according to data not shown, a *dis_steps* value of 10 allows reasonably accurate results to be produced. Each time the logical model is implemented, it must be updated *iterate_run* times to reach an attractor (steady state). At the attractor, it must be updated *iterate_record* more times in order to calculate the average activity level of each component (*Pro_i*). The Kolmogorov–Smirnov test [Xiao 2009] was used to determine these settings, which are fixed and do not require attention between studies. There is a library of synthesis rates for each of the five metabolic models. Each library entry is associated with a point in the logical model's output space. The setting, *entry_range*, indicates how close this point must lie to a set of logical model outputs for it to be a proxy of the latter. In summary, although only the values of *step*, *step_size*, and *rounds* must be determined in a study-dependent fashion, the remaining parameters are amenable to changes.
D. The calibration-related parameter are `loose' parameters which allow the multi-scale model to be fitted to experimental data. They are listed in Table S10. *Ease_restim* and *Ease_relax* indicate how sensitive an effector agent is to restimulation or the lack of it. *Ease_ACAD* and *Ease_memory* modify *Pro_ACAD* and *Pro_Mem* respectively by multiplication, thus modifying the probabilities of ACAD and memory formation. Without prior information, it was decided that *Ease_restim*, *Ease_relax*, and *Ease_ACAD* should be set to the midpoint by default, while *Ease_memory* should be an order of magnitude less by default because most effector cells die instead of turning into memory cells [Hu *et al.* 2001].

#### 5. Simulation algorithm.

At the highest level, the simulation algorithm is a series of fixed time steps, the number and size of which are determined by the duration and resolution of the modeled event, respectively. At each time step, the agents are evaluated one by one, followed by calculation of cytokine concentrations. Fig. 2b of the main manuscript illustrates the information flow between the four components of the multi-scale model. The processing and use of outputs from each module are detailed in the following paragraphs.

First, we will discuss the discrete update of each agent at each time step at the systemic/cellular scale. The most important agent behavior is its progression through the mammalian cell cycle [Harper and Brooks 2005]. In a resting state, a cell stays in the G0 phase of the cycle, but effector CD4+ T lymphocytes are metabolically different from their naive counterparts. Not only is biomass synthesis prioritized over ATP synthesis, but an effector CD4+ T lymphocyte is also not resting [MacIver *et al.* 2013]. Therefore, it begins in the G1 phase during which RNA and protein are synthesized. Subsequently, the cell moves into the S phase, where its DNA is replicated. In the G2 phase, more RNA and protein are synthesized. Finally, the cell enters the M phase to undergo mitosis and cytokinesis. When a naive agent activates and becomes an effector agent, it goes through a simplified version of this cycle. First, it enters the G1 phase to double its biomass except DNA, followed by the S phase where it doubles its DNA content. Next, the agent enters a combined G2/M phase where it divides with a probability provided by the logical model output associated with the component named `Cycle'. This description applies to reactivated memory agents too. In addition to cell cycling, each agent may perform other behaviors according to the logical model outputs, including activation (naive agents only), cell death, memory formation, migration, and cytokine secretion.

Second, we will focus on how a discrete update is performed for the logical model [Helikar *et al.* 2008, 2009, 2012, Puniya *et al.* 2018]. For each agent and in a time step at the systemic/cellular scale, multiple such updates are performed. In short, information flows from an agent and the compartment it is in to the logical model and the outputs change the agent's attributes and behaviors. Certain agent attributes and compartmental variables are used to parameterize the input components of the logical model. Then, the activity level of each non-input component is calculated and it represents the probability that the component is active given the inputs. Computationally, it means the states of all components (on or off) are updated synchronously as dictated by the logical model until convergence (estimated by the Kolmogorov-Smirnov test [Xiao 2009]). However, the nonlinear structure of the logical model can limit the maximum activity level of an output component. In other words, although each component can theoretically be 100% active [Puniya *et al.* 2018], the structure of the model may allow a maximum activity level of 70% (0.7) for a particular node only, even under optimal conditions. Therefore, the activity levels of all output components must be scaled. For example, if *Pro_Tbet* can never exceed 0.5 without scaling, but *Pro_GATA3*, *Pro_RORgt*, and *Pro_Foxp3* can reach one, then *Pro_Tbet* must be doubled before it can be compared to the other three. It is because when the logical model indicates that component Tbet is 50 % active in an agent, the cell represented by the agent actually expresses Tbet at the maximum level. In order to find the maximum activity level for each output component, 10 million random sets of inputs were generated and used to parameterize the logical model, generating 10 million sets of outputs. For each output component, the maximum activity level among the 10 million is the scale. For example, the scale of *Pro_AICD1* is 0.975, so *Pro_AICD1* must be modified by division through 0.975 to get the effective value. There is just one exception to this scaling method: the seven metabolic components. When *Pro_aGlycolysis* is 0.1, it does not mean there is a 10 % chance for aerobic glycolysis to occur. Rather, the relevant metabolic fluxes should be reduced to reflect this activity level.

The 10 million simulation results generated from the logical model were used to create an empirical distribution function for the activity level of each metabolic component. Then, a cumulative distribution function was fitted to each empirical distribution function. Physically, it indicates how unusually high an activity level is for the metabolic component in question. Numerically, it varies continuously from zero to one. For *Pro_aGlycolysis*, for example, a value of 0.2 exceeds over 90 % of the randomly generated values. According to this method, the activity level of a metabolic component is modified by applying the cumulative distribution function; the modified value (0.9 in the example about *Pro_aGlycolysis*) is utilized by the metabolic models.

Third, we will discuss how the metabolic models fit into each discrete update at the systemic/cellular scale. In short, the activity levels of the metabolic components in the logical model are used to alter the constraints of the constraint-based models, so two agents with different attributes (different activation levels, for example) may have different proliferation rates. For example, if a metabolic flux related to aerobic glycolysis has an optimized value of 1000 mmol $gDW^{-1}$ $hour^{-1}$ under optimal conditions (a transformed *Pro_aGlycolysis* of 1), the latest transformed *Pro_aGlycolysis* is 0.5, and the default constraints on this flux are -2000 and 2000 mmol $gDW^{-1}$ $hour^{-1}$, the modified constraints will be 0 and 500 mmol $gDW^{-1}$ $hour^{-1}$. The direction of the optimized flux is kept, but its value is altered to reflect the metabolic component's latest activity level. Then, under the new constraints, two objective functions are optimized. First, the synthesis rate of biomass (without DNA) is optimized for the G1 phase. Second, the synthesis rate of DNA alone is optimized for the S phase. In each case, the remaining synthesis rates (biomass without DNA, DNA, protein, ATP, and AMP) can be obtained from the optimized fluxes. It is an optimization problem which requires linear programming, a computationally expensive step. Therefore, a library of synthesis rates was generated for each of the five models in advance. This was achieved by randomly generating and transforming 10000 sets of activity levels as described in the last paragraph, and calculating the corresponding synthesis rates. At the start of a simulation, the objective functions are optimized under the default constraints for each metabolic model, leading to the optimal metabolic fluxes and thus optimal synthesis rates (biomass without DNA, DNA, protein, ATP, and AMP; both G1 and S phases). During the simulation, every time the logical model is implemented for an agent, the transformed activity levels of the metabolic components are compared to the activity levels randomly generated for the libraries. If there are close matches, the synthesis rates for the best match will be adopted. Otherwise, the logical model outputs will be used to constrain the fluxes of the metabolic model or models corresponding to the agent's phenotype, followed by optimization of the synthesis rates. In either case, if the agent assumes a mixed phenotype (*e.g.*, Th1-Th2), the corresponding libraries or metabolic models are used and the resulting synthesis rates are averaged. The updated synthesis rates are passed back to the agent, which will then updates its composition (*BM_DNA*, *DNA*, *AMP_in*, and *ATP*) accordingly. If new synthesis rates are calculated, they will be added to the libraries. In the manner of feedback control, the abundance of AMP relative to that of ATP and the synthesis rate of protein (a proxy for the abundance of ribosome [Goodsell 2000]) are converted into logical model inputs.

#### 6. MATLAB program used to implement the multi-scale model in order to simulate the evolution of agents and cytokine concentrations in response to an input signal.

The multi-scale model is stochastic because of the logical model and most agent behaviors, so it must be implemented multiple times in order to obtain a distribution of results (Monte Carlo method). In other words, the MATLAB program whose pseudocode is as follows must be run multiple times.

1. Populate the compartments with naive agents.
2. Initialize the cytokine concentrations in the compartments.
3. At the beginning of each time step, evaluate each agent.
    a. If *State* is 0, it is a dead agent, so ignore it.
    b. If *State* is 1, it is a naive agent.
        i. If TCR signaling is on in the lymphoid tissues according to $TCR_{t,2}$, boost its migration thither.
        ii. If TCR signaling is on in the lymphoid tissues according to $TCR_{t,2}$ and it is in the lymphoid tissues, stimulate it once by incrementing *TCR_stimuli* by one, *TCR_strength* by $TCR_{t,2}$, *CD28_strength* by $CD28_{t,2}$, and *i_sum* by $C_{2,i}$.
        iii. If $TCR\_stimuli = act\_thres$, change *State* to 2, implement the logical model, and update the synthesis rates according to the constraint-based models.
    c. If *State* is 2, it is an effector agent.
        i. Change its migration scheme so that it travels unidirectionally from the lymphoid tissues to the target organ via the circulatory system.
        ii. Implement the logical model, update the synthesis rates according to the constraint-based models, and update its composition, hence deciding its *Phase*.
        iii. If $Phase = 3$ and $Divisions < div\_lim$, it divides into two agents stochastically according to *Pro_Cycle*. After division, each daughter agent returns to the G1 phase ($Phase = 1$). Update *Resist* and *Plasticity*, implement the logical model, and update the synthesis rates according to the constraint-based models. Also, if $Divisions \geq mig\_thres$, update its migration scheme so that it will leave the lymphoid tissues for the circulatory system.
        iv. If TCR signaling is on according to $TCR_{t,i}$, it may undergo AICD.

v. If TCR signaling is off according to $TCR_{t,i}$, it may become a memory agent. If it does, *State* becomes 3 and *act_thres* is incremented by *act_thres_mem*. If it does not, it may undergo ACAD. If neither happens, its activation level in terms of *TCR_strength* and *CD28_strength* will decrease.

d. If *State* is 3, it is a memory agent.

   i. If TCR signaling is on according to $TCR_{t,i}$, stimulate it once by incrementing *TCR_stimuli* by one.

   ii. If $TCR\_stimuli = act\_thres$, change *State* to 4, set *TCR_strength* and *CD28_strength* to *TCR_max* and *CD28_max* respectively. Then, update the migration scheme so that it travels unidirectionally from the lymphoid tissues to the target organ via the circulatory system. Finally, implement the logical model and update the synthesis rates according to the constraint-based models.

e. If *State* is 4, it is a reactivated memory agent, which behaves in exactly the same way as an effector agent. Therefore, the same computational steps apply.

4. At the same time step, redistribute every agent between the three compartments according to its *k_tot*, *k_toln*, and *k_toc*. This step applies to the agents created at this time step too.
5. At the same time step, after redistribution, check the number of naive agents in each compartment. When the population is below the threshold defined by *Refill_thres*, add new naive agents to restore their numbers to $N_{t0}$, $N_{ln0}$, and $N_{c0}$.
6. At the end of the time step, calculate the cytokine concentrations in the three compartments by solving their governing ordinary differential equations. The agents created at this time step do not count towards this calculation.
7. Parameterize the ordinary differential equations for the next time step. For each agent whose *State* is 2 or 4, evaluate its ability to secrete cytokines (IL2, IL4, IL6, IL17, IL21, and IFNg) according to the relevant attributes such as *Pro_IL2*. If *Plasticity* is high, the agent has not divided many times and secretes IL2 only. If *Plasticity* is low, it is highly differentiated and can potentially secrete all six cytokines.
8. Return to step 3 and begin the next time step.

#### 7. Default input signal.

The viral trajectory from an experimental study about influenza [Román *et al.* 2002] was used as the input signal in all presented simulations except for those of hypothetical infections. In that study, mice were inoculated intranasally with the influenza A virus (A/PR/8/34). After that, viral titer in their infected lungs and the population sizes of CD4+ T cells in different tissues were determined.

In that paper, the measurements are presented on a logarithmic scale. In the multi-scale model, however, the signal is non-physical, so the data was abstracted into a signal on a linear scale between zero and one. When the input signal is one, TCR signaling is deterministically on. Therefore, the maximum was not set to one to allow some stochastic effects in the whole time window. That aside, 0.8 was chosen arbitrarily. For the sake of simplicity, the experimentally observed decline between two and six days post-infection was ignored. Although measurements were taken up to 10 days post-infection in the experiment, an extended time frame (until 400 hours post-infection) was chosen to ensure a full reading in the presented simulations. The overall temporal profile of this signal is shown in Fig. 5a in the main manuscript (trajectory labeled 0.8).

When the relevant simulations were set up, this default input signal was applied in the target organ (lungs) and lymphoid tissues (draining lymph node). In the circulatory system, an always-off signal was chosen due to the assumption of a negligible viral concentration. This assumption is reasonable because the circulatory system is far bigger than the other two compartments.

#### 8. Input signals representing hypothetical infections.

The input signals for five hypothetical acute infections by the influenza A virus, alongside the default input signal (for an actual acute infection by the influenza A virus), are shown in Fig. 5a of the main manuscript. Each of them differs from the default in three ways. First, the peak is not at 0.8. Second, the rise from zero to the said peak in the first two days post-infection has a different rate; it is steeper if the peak is above 0.8, flatter if below; what is conserved is the linear nature of the rise. Third, the drop from the peak to zero between five and 10 days post-infection has a different rate; it is steeper if the peak is above 0.8, flatter if below; what is conserved is the linear nature of the drop.

The input signal for a hypothetical chronic infection by the influenza A virus is shown in Fig. 5e of the main manuscript. It differs from the default input signal in its pattern. Like the default input signal, it rises linearly from zero to 0.8 in the first two days. However, unlike the default, this chronic input signal does not fall after reaching 0.8. Furthermore, the duration of this signal is 10 times longer than that of the default, 4000 hours as opposed to 400 hours.

#### 9. Details on the cytokine combinations and dosages used in the presented simulations.

The cytokine combinations and dosages used in the simulations presented in the main manuscript were also used in an experimental study [Eizenberg-Magar *et al.* 2017].

In that study, CD4+ T lymphocytes were isolated from mouse spleens by magnetic microbeads and cultured with plate-bound anti-CD3 and anti-CD28 agents. The resulting cell cultures were supplemented with different binary combinations of IL2 (5 ng/mL), IL4 (20 ng/mL), IL6 (20 ng/mL), IL12 (10 ng/mL), IFNg (10 ng/mL), and TGFβ (10 ng/mL). It means that each cytokine was either present at the indicated concentration or absent. After four days of stimulation, each culture was transferred to an uncoated plate where it stayed for three more days. After that, each culture medium was washed and the cells were transferred to a new plate, coated with anti-CD3 and anti-CD28 agents, bathed in a fresh medium with the same cytokines. After four days, the culture was transferred to an uncoated plate where it stayed for three more days. The expression levels of cytokines and transcription factors were measured by intracellular antibody staining and flow cytometry.

The results indicate stereotypical expression patterns. For example, in the culture with IL2 and IL4, the cells strongly expressed several Th2-associated cytokines (IL4, IL5, and IL12) and the Th2-defining transcription factor (GATA3). In conclusion, IL12 led to the Th1 phenotype; IL2 and IL4, Th2; IL6 and TGFβ, Th17; and IL2 and TGFβ, Treg.

#### 10. Convergence study.

An input signal was designed to mimic the viral dynamics observed in an influenza study [Román *et al.* 2002]; the temporal dynamics of this signal are shown in Fig. 5a of the main manuscript (trajectory labeled 0.8); the details are in the section dedicated to the default input signal.

The initial cytokine concentrations were set to the values used in a different experimental study [Eizenberg-Magar *et al.* 2017], wherein only IL2, IL4, IL6, IL12, IFNg, and TGFβ were used. A summary of this study can be found in the section describing and explaining the cytokine concentrations used in the simulations presented in the main manuscript. The cytokine production terms associated with the input signal were set to zero.

A Monte Carlo simulation was run to simulate the population dynamics of CD4+ T cells in response to the default signal. In this simulation, the multi-scale model was implemented 100 times to generate 100 realizations of the modeled system. For each realization, the maximum number of effector and reactivated memory cells (metric 1) and the final number of memory cells (metric 2) were recorded. 100 data points were thus obtained for each metric: one for each realization. For each metric, the 100 data points were resampled with replacement to generate batches of different sizes (10 to 500 data points); an average was then calculated for each batch. 50 averages were calculated for each metric in this manner.

Based on the results (Fig. S2 and Fig. S3), 50 realizations ( $rounds = 50$ ) are sufficient for obtaining robust average results. As can be seen in each figure, the fluctuations of the average metric between 50 and 500 realizations are less than 7 % of the average metric obtained from all 500 realizations. Although an increase in *rounds* can increase the robustness further, the extra computational cost is not justified.

#### 11. Estimating logical model attractors corresponding to different cytokine combinations and dosages.

Fig. 4a in the main manuscript presents the transformed activity levels of selected components of the logical model in the estimates of five attractors corresponding to five cytokine combinations. They are estimates because in the presence of stochastic inputs, it is not feasible to calculate attractors *per se*. Each estimate was computed as follows. The first class of inputs (*IL2_e*, *IL4_e*, *IL6_e*, *IL12*, *IL18*, *IL23*, *IL27*, *IFNg_e*, and *TGFβ*) was informed by the cytokine concentrations used in an experimental study [Eizenberg-Magar *et al.* 2017], as explained in a previous section. For the second class, both *TCR* and *CD28* were set to 0.5, the unbiased midpoint. The same reason was behind the decision to set *Ribosome* and *Resist* in the third class to 0.5, but *mTORC1_t* and *AMP* were set to 0.05. This reflects the prioritization of biomass synthesis over energy generation. Indeed, since the estimates were computed for effector cells, the energy-generating pathways were not of interest, so *mTORC1_t* and *AMP* were set at a low level. With these input conditions, the logical model was implemented as detailed in the section about the simulation algorithm.

For each of the five cytokine combinations, 10000 simulations were run, and the 10000 estimated attractors were averaged to give the results in Fig. 4a. We found that running more than 10000 simulations did not change the activity levels significantly for the case with all six cytokines. As an example, averaging 100000 simulations gave *Tbet* as 0.1537; 10000 simulations, 0.1545; 1000 simulations, 0.1543; 100 simulations, 0.1514; and 10 simulations, 0.1670. We acknowledge that the average for each cytokine combination hides a distribution of estimated attractors corresponding to that combination, so the results in Fig. 4a should be interpreted as typical behaviors.

The estimated attractors presented in Fig. 4b in the main manuscript were computed similarly. In each of the three cases, *IL4_e*, *IL6_e*, *IL12*, *IL18*, *IL23*, *IL27*, *IFNg_e*, and *TGFβ* were set to zero. For the reasons provided above, *TCR*, *CD28*, *Ribosome*, and *Resist* were set to 0.5; *mTORC1_t* and *AMP* to 0.05. In the three cases, *IL2_e* was set to match 0, 5, and 50 ng/mL respectively.

#### 12. Re-using the multi-scale model to study diverse immune phenomena by changing the parameters, running Monte Carlo simulations, and analyzing the distribution of results.

The multi-scale model has four classes of variables and parameters. In the first three classes, most of the parameters are fixed and based on findings in the literature, but some must be tailored for the problem of interest. The fourth class has default values, but they can be changed in order to fit the multi-scale model to experimental data.

The following steps represent a concise summary of the process. In essence, parameter values that are appropriate for the study must be decided upon and used. For example, if it is about influenza, the target organ will be the lungs, so the latter's volume will be used for $V_1$. If experimental datasets are available, the calibration parameters can be used for model fitting.

1. Set up the compartments by providing $V_1$, $V_2$, $V_3$, $Q_a$, and $Q_b$.
2. Populate the compartments by fixing the numbers of agents in them: $N_1$, $N_2$, $N_3$, $N_{1,p}$, $N_{2,p}$, and $N_{3,p}$. Computational resources are saved if only a fraction of the cell population is simulated; it is equivalent to simulating a portion of each compartment. For example, if $N_1$ is half of $N_{1,p}$, the target organ compartment will have an effective volume of $\frac{V_1}{2}$, a fact accounted for when cytokine concentrations are calculated.
3. Introduce cytokines to the compartments by providing their initial cytokine concentrations ($C_{i,j,0}$).
4. Provide the temporal trajectory of the input signal ($s_{t,i}$), as well as the cytokine production rates associated with it ($P_{i,j}^{in}$).
5. Set up the numerical methods by providing the number of time steps (*steps*), the step size (*step_size*), and the number of times the model must be implemented for convergence (*rounds*), as well as deciding whether to use the default values of the remaining parameters in Table S9.
6. Decide on the four calibration parameters in relation to the problem under consideration and any experimental data available for calibration.

Although the remaining parameters are considered fixed, they can be changed in order to experiment with the intrinsic properties of CD4+ T cells. For example, a smaller value of *act_thres_naive* means a more responsive population of naive cells.

After the multi-scale model has been parameterized and initialized for a particular problem, a convergence study must be carried out. The multi-scale model is stochastic in many aspects such as the use of a stochastic logical model; as such, it must be implemented multiple times in order to obtain robust average results (Monte Carlo simulation). The number of times it must be implemented to achieve convergence is the value of *rounds*.

### Figure S1

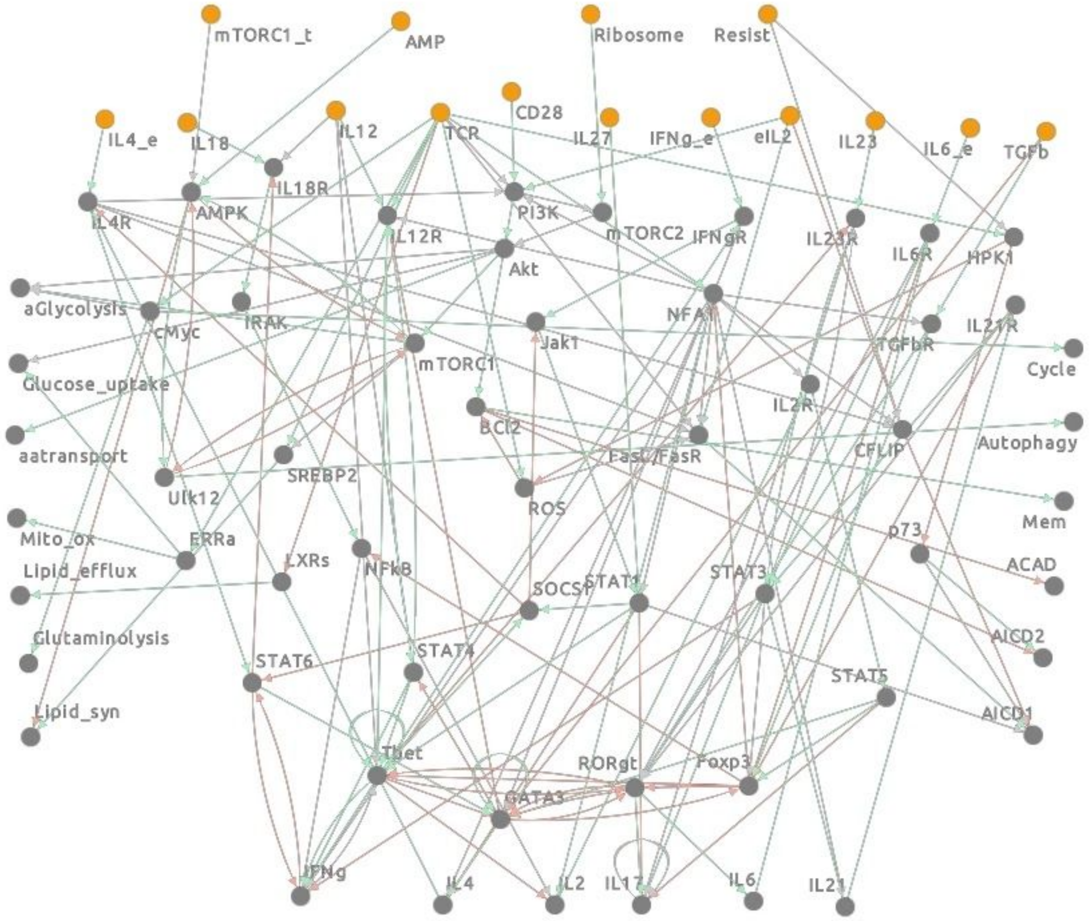


Expanded version of a previously published logical model [Puniya *et al.* 2018]. The yellow nodes are input components, while the grey ones include the output components which influence other modules (such as metabolic models) of the multi-scale model and the internal components which only influence the logical model itself. Each green arrow represents an activating interaction, while each red arrow represents an inhibiting interaction.

### Figure S2

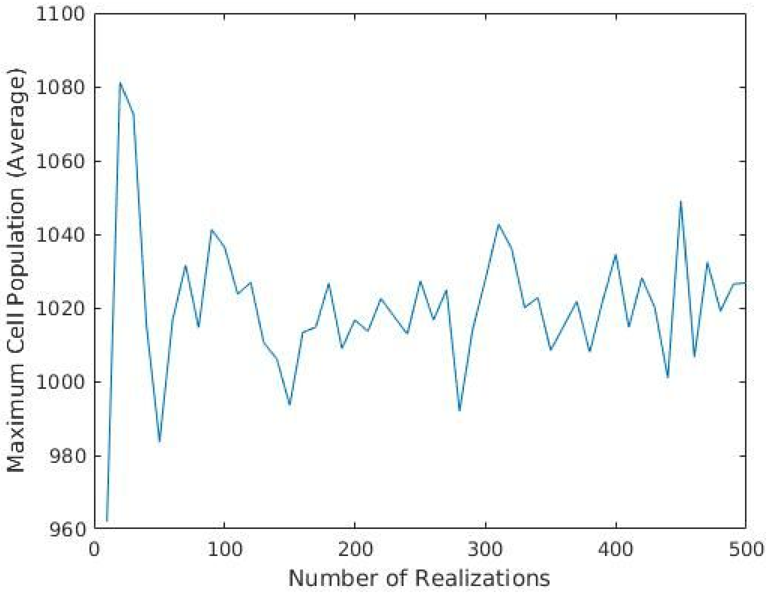


Convergence plot for the maximum population of CD4+ T lymphocytes (effector and reactivated memory cells only) in an influenza infection. 100 realizations of the modeled system were produced in a Monte Carlo simulation. The maximum population size was calculated for each realization. Then, batches of these quantities were produced by sampling with replacement. Each batch has a different size, so the x-axis describes the number of realizations. For each batch, the average was calculated and plotted on the y-axis.

Figure S3

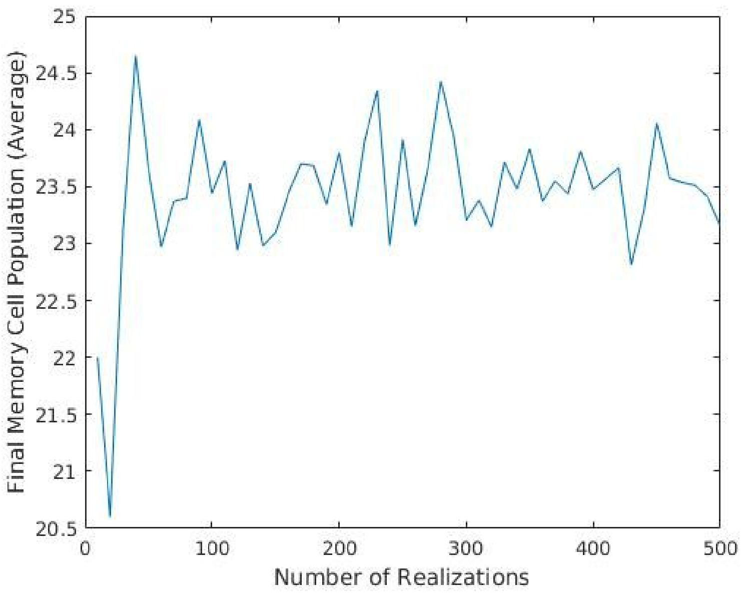


Convergence plot for the final number of memory CD4+ T lymphocytes in an influenza infection. 100 realizations of the modeled system were produced in a Monte Carlo simulation. The final memory cell count was calculated for each realization. Then, batches of these quantities were produced by sampling with replacement. Each batch has a different size, so the x-axis describes the number of realizations. For each batch, the average was calculated and plotted on the y-axis.

### Table S1

Attributes/variables that define the metabolic state of an agent; G1 and S refer to the first two phases in the cell cycle the agent goes through; the third and final one is G2/M.

| Variable | Description | Units |
|---|---|---|
| *BM_DNA* | Amount of biomass minus DNA | g |
| *DNA* | Amount of DNA | g |
| *AMP_in* | Concentration of AMP | M |
| *ATP* | Concentration of ATP | M |
| *G1_BM_DNA* | Synthesis rate of biomass minus DNA during the G1 phase | $g \cdot h^{-1}$ |
| *G1_DNA* | Synthesis rate of DNA during the G1 phase | $g \cdot h^{-1}$ |
| *G1_AMP* | Synthesis rate of AMP during the G1 phase | $M \cdot h^{-1}$ |
| *G1_ATP* | Synthesis rate of ATP during the G1 phase | $M \cdot h^{-1}$ |
| *S_BM_DNA* | Synthesis rate of biomass minus DNA during the S phase | $g \cdot h^{-1}$ |
| *S_DNA* | Synthesis rate of DNA during the S phase | $g \cdot h^{-1}$ |
| *S_AMP* | Synthesis rate of AMP during the S phase | $M \cdot h^{-1}$ |
| *S_ATP* | Synthesis rate of ATP during the S phase | $M \cdot h^{-1}$ |

### Table S2

Parameters for the metabolizing behaviors of an agent; it takes in metabolites and converts them into different kinds of products in different phases of the cell cycle; at the end of the cycle, it divides into two identical agents. If patient-specific and single-cell data are available, these parameters can be changed to absorb the data.

| Parameter | Description | Value | Units | References |
|---|---|---|---|---|
| *BM_DNA_initial* | Amount of biomass minus DNA at the start of a cell cycle | $5.65 \times 10^{-11}$ | g | [Segel *et al.* 1981, Cooper *et al.* 2000, Gillooly *et al.* 2015] |
| *DNA_initial* | Amount of DNA at the start of a cell cycle | $8 \times 10^{-12}$ | g | [Gillooly *et al.* 2015] |
| *AMP_initial* | Concentration of AMP at the start of a cell cycle | $8.5 \times 10^{-8}$ | M | [Hardie and Hawley 2001] |
| *ATP_initial* | Concentration of ATP at the start of a cell cycle | $8.5 \times 10^{-6}$ | M | [Mikirova *et al.* 2005] |
| *cell_volume* | Cell volume | $5.236 \times 10^{-13}$ | $dm^3$ | [Anaya *et al.* 2013] |

### Table S3

Attributes/variables used to parameterize the logical model or determined by its outputs. The signaling strengths are fixed by the input signal, *i_sum*'s by the cytokine concentrations throughout the activation stage, and the rest by the internal state of the agent for which the logical model is implemented.

| Variable | Description | Units |
|---|---|---|
| *TCR_strength* | TCR signaling strength in the agent | None |
| *CD28_strength* | CD28 signaling strength in the agent | None |
| *i_sum* | Abundance of cytokine *i* throughout the activation stage (nine in total: IL2, IL4, IL6, IL12, IL18, IL23, IL27, IFNg, and TGFβ) | M |
| *Resist* | Probability of resistance to apoptosis | None |
| *Ribosome* | Abundance of ribosome (zero to one, continuous) | None |
| *mTORC1_t_count* | Number of times the logical model has been implemented and the outcome has led to mTORC1 expression in the agent | None |
| *BN_count* | Number of times the logical model has been implemented for the agent | None |
| *Pro_i* | Activity level of component *i* or the probability that it is on at the attractor following an implementation of the logical model (58 in total) | None |

### Table S4

Parameters used to turn certain agent attributes into inputs for the logical model. The parameter *step_size* is the size of each time step in hours; it is the time interval of each update when the multi-scale model is implemented. The parameters about cytokine receptors can absorb patient-specific and cytokine-specific data, if available; the listed values are based on experiments on IL2.

| Parameter | Description | Value | Units | References |
|---|---|---|---|---|
| *TCR_max* | Maximum TCR signaling strength | $\frac{24}{step\_size}$ | None | [Jelley-Gibbs *et al.* 2000] |
| *CD28_max* | Maximum CD28 signaling strength | $\frac{24}{step\_size}$ | None | [Jelley-Gibbs *et al.* 2000] |
| *rep_min* | Minimum number of receptors for each cytokine type expressed by the agent | 100 | None | [Foxwell *et al.* 1992, McKinstry *et al.* 1997] |
| *rep_max* | Maximum number of receptors for each cytokine type expressed by the agent | 6000 | None | [Foxwell *et al.* 1992, McKinstry *et al.* 1997] |
| *sat_count* | Minimum number of cytokine-receptor complexes needed for peak downstream signaling in the agent | 1500 | None | [Fallon and Lauffenburger 2000] |
| *BN_count_thres* | Minimum number of logical model implementations before the persistence of mTORC1 expression is important in the agent | 10 | None | Assumed |

### Table S5

Attributes/variables internal to an agent in the sense that they do not directly influence the other constituent models of the multi-scale model.

| Variable | Description | Units |
|---|---|---|
| *State* | Agent state (0 = dead, 1 = naive, 2 = effector, 3 = memory, and 4 = reactivated memory) | None |
| *Phenotype* | Th0, Th1, Th2, Th17, Treg, and every mixed state | None |
| *Phase* | Cell cycle phase (1 = G1, 2 = S, and 3 = G2/M) | None |
| *Location* | Compartment the agent is in (1 = target organ, 2 = lymphoid tissues, and 3= circulatory system) | None |
| *TCR_stimuli* | Number of stimuli accumulated by the agent | None |
| *act_thres* | Number of stimuli needed for activation or reactivation | None |
| *Divisions* | Number of division rounds the agent has undergone | None |
| *Plasticity* | Probability of transdifferentiation | None |
| *k_tot* | Probability of migration to the target organ | None |
| *k_toln* | Probability of migration to the lymphoid tissues | None |
| *k_toc* | Probability of migration to the circulatory system | None |

### Table S6

Parameters that define an agent and the agent population. The parameter *step_size* is the size of each time step in hours; it is the time interval of each update when the multi-scale model is implemented. Patient-specific data can be integrated into the model through the cell counts and *k_effector_CtoT*. Patient-specific and cytokine-specific data can be used to optimize $P_{ex}$.

| Parameter | Description | Value | Units | References |
|---|---|---|---|---|
| *ID* | Agent identifying number | Assigned | None | Assigned |
| $N_{t0}$ | Initial number of naive agents in the target organ compartment | Study-dependent (32 for the lungs) | None | [Bui *et al.* 2007, Ganusov and De Boer 2007, Purwar *et al.* 2011, Gasper *et al.* 2014] |
| $N_{ln0}$ | Initial number of naive agents in the lymphoid tissue compartment | Study-dependent (10 for one draining lymph node) | None | [Bui *et al.* 2007, Ganusov and De Boer 2007, Gasper *et al.* 2014, Margaris and Black 2012, Sathaliyawala *et al.* 2013] |
| $N_{c0}$ | Initial number of naive agents in the circulatory system compartment | 391 | None | [Bui *et al.* 2007, Ganusov and De Boer 2007, Gasper *et al.* 2014, Sathaliyawala *et al.* 2013] |
| $N_{t0\_all}$ | Initial number of naive CD4+ T cells in the actual target organ | Study-dependent ($1.48 \times 10^6$ for the lungs) | None | [Bui *et al.* 2007, Ganusov and De Boer 2007, Purwar *et al.* 2011, Gasper *et al.* 2014] |
| $N_{ln0\_all}$ | Initial number of naive CD4+ T cells in the actual lymphoid tissues | Study-dependent ($4.56 \times 10^5$ for one draining lymph node) | None | [Bui *et al.* 2007, Ganusov and De Boer 2007, Gasper *et al.* 2014, Margaris and Black 2012, Sathaliyawala *et al.* 2013] |

| | | | | |
|---|---|---|---|---|
| $N_{c0\_all}$ | Initial number of naive CD4+ T cells in the actual circulatory system | $1.785 \times 10^7$ | None | [Bui *et al.* 2007, Ganusov and De Boer 2007, Gasper *et al.* 2014, Sathaliyawala *et al.* 2013] |
| *Refill_thres* | At this fraction of the initial population, new naive agents are added to the compartments | 0.2 | None | Assumed |
| *act_thres_naive* | Number of stimuli the agent needs before it can move from the naive state to the effector state | $\frac{24}{step\_size}$ | None | [Jelley-Gibbs *et al.* 2000] |
| *act_thres_mem* | Number of stimuli the agent needs before it can move from the memory state to the reactivated memory state | $\frac{2}{step\_size}$ | None | [Bevington *et al.* 2017] |
| *div_lim* | Maximum number of division rounds the agent can undergo | 10 | None | [Parkin and Cohen 2001] |
| *mig_thres* | Number of division rounds the agent must undergo before it can exit the lymphoid tissues | 6 | None | [Román *et al.* 2002] |
| *k_toln_boost* | When there is an infection and the agent is in the naive state, its rate of migration from the target organ to the lymphoid tissues increases by this factor | 5 | None | [Soderberg *et al.* 2005] |
| *k_effector_CtoT* | Probability that the agent will migrate to the target organ when it is in the effector state and in the circulatory system | $0.02083 \times step\_size$ | None | [McKinstry *et al.* 2010] |
| $P_{ex}$ | Rate of cytokine production by the agent (not cytokine-specific) | 10 | Molecules·$s^{-1}$ | [Han *et al.* 2012] |

### Table S7

Variables that describe the changing aspects of the compartments. Although $s_{t,\,i}$ was pre-defined in the presented simulations, it can be replaced by a time-dependent function of other variables; it is study-dependent and can absorb patient-specific data. Although $TCR_{t,\,i}$ and $CD28_{t,\,i}$ were equated to $s_{t,\,i}$ in the presented simulations, they can be replaced by functions; they are study-dependent and can absorb patient-specific data. Overall, these three variables can be tailored in order to study different infections in different patients.

| Variable | Description | Units |
|---|---|---|
| $s_{t,\,i}$ | Input signal in compartment *i* at time *t* (continuous signal between zero and one); defined in a study-dependent manner | None |
| $TCR_{t,\,i}$ | TCR signaling strength in compartment *i* at time *t* (continuous signal between zero and one); defined in a study-dependent manner (same as $s_{t,\,i}$ by default) | None |
| $CD28_{t,\,i}$ | CD28 signaling strength in compartment *i* at time *t* (continuous signal between zero and one); defined in a study-dependent manner (same as $s_{t,\,i}$ by default) | None |
| $C_{i,j}$ | Concentration of cytokine *j* in compartment *i* (11 cytokines in total: IL2, IL4, IL6, IL12, IL17, IL18, IL21, IL23, IL27, IFNγ, and TGFβ) | M |

### Table S8

Parameters that define the unchanging aspects of the compartments. The compartment volumes and intercompartmental flow rates allow patient-specific data to be integrated into the model. The cytokine production rates associated with the input signal, as well as the initial cytokine concentrations, are study-dependent and can absorb patient-specific data. The cytokine production rates should be defined in conjunction with the input signal.

| Parameter | Description | Value | Units | References |
|---|---|---|---|---|
| $V_1$ | Volume of the target organ | Study-dependent (0.547 for the lungs) | $dm^3$ | [Ryman and Meibohm 2017] |
| $V_2$ | Volume of the lymphoid tissues | Study-dependent ($4.545 \times 10^{-4}$ for one draining lymph node) | $dm^3$ | [Genereux and Howie 1984, Jensen *et al.* 2010] |
| $V_3$ | Volume of the circulatory system | 3.8 | $dm^3$ | [Ryman and Meibohm 2017] |
| $Q_a$ | Flow rate from the circulatory system to the target organ | Study-dependent (356 for the lungs) | $dm^3 \cdot h^{-1}$ | [Ryman and Meibohm 2017] |
| $Q_b$ | Flow rate from the target organ to the lymphoid tissues | Study-dependent (0.0094 for the lungs and its nearest draining lymph node) | $dm^3 \cdot h^{-1}$ | [Ryman and Meibohm 2017] |
| $k_{i,j}^{deg}$ | Degradation rate constant of cytokine $j$ in compartment $i$ | 0.1 | $h^{-1}$ | [Su *et al.* 2009] |
| $\tau$ | Time scale | Size of a time step (*step_size*) | h | Study-dependent |
| $C_{i,j,0}$ | Initial concentration of cytokine $j$ in compartment $i$ | Study-dependent | M | Study-dependent |
| $C_{j,s}$ | Concentration scale of cytokine $j$ | $1 \times 10^{-8}$ | M | [Burska *et al.* 2014] |
| $P_{i,j}^{in}$ | Production rate of cytokine $j$ in compartment $i$ due to the input signal | Study-dependent | $M \cdot h^{-1}$ | Study-dependent |

### Table S9

Parameters for the numerical methods used to implement the multi-scale model. The values of *steps* and *step_size* are defined when the input signal is defined; they depend on the duration of the modeled infection and the time resolution required by the study respectively. A convergence study is needed to fix *rounds* after the multi-scale model is parameterized for a specific study.

| Parameter | Description | Value | Units | References |
|---|---|---|---|---|
| *steps* | Number of time steps | Study-dependent | None | Study-dependent |
| *step_size* | Size of each time step | Study-dependent (one by default) | hours | Study-dependent |
| *rounds* | Number of times the multi-scale model must be implemented | Study-dependent | None | Study-dependent |
| *dis_steps* | Number of discrete steps the ode45 solver must take | Study-dependent (10 by default) | None | Study-dependent |
| *iterate_run* | Number of discrete updates before the logical model converges | 250 | None | Kolmogorov–Smirnov test [Xiao 2009] |
| *iterate_record* | Number of discrete updates needed to sample an attractor of the logical model | 200 | None | Kolmogorov–Smirnov test [Xiao 2009] |
| *entry_range* | Search range in the library of synthesis rates (zero to one, continuous) | Study-dependent (0.05 by default) | None | Study-dependent |

### Table S10

Parameters for model calibration. These are `loose' parameters which can be varied to calibrate the multi-scale model for a specific study. For example, if the CD4+ T cell dynamics are known for an influenza infection, a grid search can be performed: implement the model with different parametric combinations, compare the simulation results with the known dynamics, and pick the parametric combination for the closest match.

| Parameter | Description | Value | Units | References |
|---|---|---|---|---|
| *Ease_restim* | Ease of restimulation | Study-dependent (0.5 by default; zero to one, continuous) | None | Study-dependent |
| *Ease_ACAD* | Ease of ACAD | Study-dependent (0.5 by default; zero to one, continuous) | None | Study-dependent |
| *Ease_memory* | Ease of memory formation | Study-dependent (0.05 by default; zero to one, continuous) | None | Study-dependent |
| *Ease_relax* | Ease of relaxation | Study-dependent (0.5 by default; zero to one, continuous) | None | Study-dependent |
